# IGF2BP3 amplifies antiviral innate immunity with implications for autoimmune diseases

**DOI:** 10.64898/2026.08.01.742171

**Authors:** Ao Zhang, Shijin Geng, Rong-Chun Tang, Hengxiang Yu, Yunxuan Zhou, Lan Zhang, Xiuyuan Sun, Jun Zhang

## Abstract

The insulin-like growth factor 2 mRNA-binding protein 3 (IGF2BP3) is a known N6-methyladenosine (m^6^A) reader, but its role in antiviral innate immunity is unknown. Here, we identify IGF2BP3 as a critical positive regulator of antiviral responses. Viral infection and interferon (IFN) stimulation upregulate IGF2BP3, establishing a feedforward loop that potentiates virus-induced activation of the TBK1–IRF3 and NF-κB pathways, thereby amplifying type I interferon (IFN-I) production, and restricting viral replication in human and murine cells and *in vivo*. Mechanistically, IGF2BP3 directly binds and stabilizes *MAVS* and *TBK1* mRNAs and promotes their translation by facilitating recruitment to the eIF4F/PABP-associated initiation complex. Upon infection, IGF2BP3 relocalizes to antiviral stress granules (avSGs), where it scaffolds the RIG-I–G3BP1 complex to enhance viral RNA sensing. Notably, IGF2BP3 is aberrantly upregulated in patients with systemic lupus erythematosus (SLE) and in *Trex1* knockout (KO) mice, and pharmacological inhibition by curcumol suppresses IFN-I–driven pathology and improves survival. Collectively, our findings establish IGF2BP3 as a central feedforward circuit that couples viral RNA sensing to the control of RNA stability and translation of key signaling molecules, and reveals its potential as a therapeutic target in interferon-associated autoimmune diseases.

**Significance Statement:** Antiviral immunity demands rapid and coordinated gene expression, yet how RNA-binding proteins link viral recognition to downstream signaling is poorly understood. We reveal that the m^6^A reader IGF2BP3 acts as a central amplifier of antiviral innate immunity. Induced by both viruses and interferons, IGF2BP3 enhances viral RNA sensing, stabilizes key signaling transcripts, and boosts their translation, creating a self-reinforcing feedforward circuit that strengthens interferon responses. Beyond host defense, IGF2BP3 is aberrantly elevated in interferon-driven autoimmunity, and its pharmacological inhibition reduce disease pathology *in vivo*. Our findings uncover a previously unrecognized mechanism that integrates RNA metabolism, stress-granule signaling, and translational control to regulate innate immunity, offering new therapeutic perspectives for both infectious and autoimmune disease.

## Introduction

The innate immune system serves as the fundamental defense mechanism against viral pathogens, relying on pattern recognition receptors (PRRs) to detect pathogen-associated molecular patterns (PAMPs) and trigger antiviral responses via interferon signaling pathways and inflammatory cytokine production(1–3). The precise regulation of these responses is essential for effective viral clearance while preventing excessive immune activation that can cause tissue damage or autoimmune disorders(4, 5). Emerging evidence has highlighted a pivotal role of post-transcriptional modifications, particularly m^6^A, in modulating immune-related gene expression and antiviral defense(6, 7). As the most abundant internal mRNA modification in eukaryotic cells, m^6^A dynamically regulates RNA metabolism, including splicing, stability, and translation efficiency, through recognition by specific RNA-binding proteins termed "readers"(8–10). However, the molecular mechanisms by which distinct m^6^A readers interpret this epitranscriptomic mark to control antiviral innate immunity remain incompletely understood.

Among the m^6^A-binding proteins, IGF2BP family (IGF2BP1,2,3) has emerged as a critical regulator of RNA fate(11, 12). Unlike the well-characterized YTH domain-containing family proteins (YTHDF1,2,3 and YTHDC1,2), which direct multiple steps of mRNA fate(13–15), IGF2BP proteins constitute a distinct class of m^6^A readers that preferentially stabilize target transcripts and, in certain contexts, enhance their translation, often in concert with additional RNA-binding proteins(11). Current understanding of IGF2BP3’s biological functions is primarily in cancer, where it promotes tumor cell proliferation, invasion, and metastasis by stabilization mRNAs encoding growth factors and cell cycle regulators(16–18). However, accumulating evidence points to broader physiological roles. Notably, recent studies have demonstrated that IGF2BP3 modulates immune cell function, including macrophage polarization, by stabilizing transcripts involved in inflammatory signaling pathways(19, 20). Considering that m^6^A modification is dynamically regulated during viral infection and profoundly influences host-pathogen interactions(21, 22), we hypothesized that IGF2BP3 may serve as a critical epitranscriptomic regulator of antiviral immune responses.

In parallel, avSGs have been recognized as signaling hubs that concentrate RNA-binding proteins and facilitate RIG-I–mediated viral RNA sensing, raising the possibility that m^6^A readers may influence RNA detection and downstream signaling within these condensates(23–25). Moreover, because IFN-I circuits that are protective during acute infection can become pathogenic when chronically activated virus-inducible amplifiers of IFN-I signaling could plausibly contribute to interferon-associated autoimmune inflammation(26–28).

In this study, we systematically investigate the role of IGF2BP3 in modulating innate immune responses to viral infection. Through comprehensive approaches, we demonstrate that IGF2BP3 selectively stabilizes key antiviral transcripts and facilitates their translation, and it additionally associates with antiviral SG components to facilitate RIG-I–dependent viral RNA sensing, thereby amplifying the IFN-I response. Our findings uncover a novel regulatory mechanism by which IGF2BP3-mediated RNA stabilization and translational enhancement fine-tunes antiviral defense, providing important insights into the complex interplay between epitranscriptomic regulation and innate immunity. These results not only expand our understanding of m^6^A reader proteins in immune regulation but also suggest potential therapeutic strategies for modulating interferon-associated autoimmune inflammation through targeting IGF2BP3.

## Results

### IGF2BP3 is a virus-inducible m6A reader that amplifies antiviral signaling

To systematically assess whether viral infection alters the expression of m^6^A-related regulators, we analyzed transcriptomic datasets from patients with influenza (GSE100160) and COVID-19 (GSE217948). Among key m^6^A regulators, *IGF2BP3* was consistently and markedly upregulated in both cohorts, whereas changes in most other regulators were limited (Supplemental Figure 1, A–P). Notably, *IGF2BP3* expression positively correlated with IFN-I genes, and Gene Ontology analysis (GO) of IGF2BP3-correlated genes revealed enrichment for antiviral and IFN-I signaling pathways (Supplemental Figure 1, Q and R), suggesting a potential role for IGF2BP3 in antiviral immunity.

We next validated IGF2BP3 induction experimentally by different stimuli. In THP-1 cells and immortalized bone marrow–derived macrophages (iBMDMs), Sendai virus (SeV) infection or IFN-α/β stimulation increased both *IGF2BP3* mRNA and IGF2BP3 protein levels (Figure 1, A–C). This induction was not restricted to RNA viruses, as HSV-1 infection also upregulated *IGF2BP3* mRNA in multiple cell types (Supplemental Figure 1S). Moreover, LPS stimulation elevated *IGF2BP3* expression in THP-1 cells and primary BMDMs (Supplemental Figure 1T), indicating that IGF2BP3 is broadly responsive to innate immune activation.

**Figure 1.**
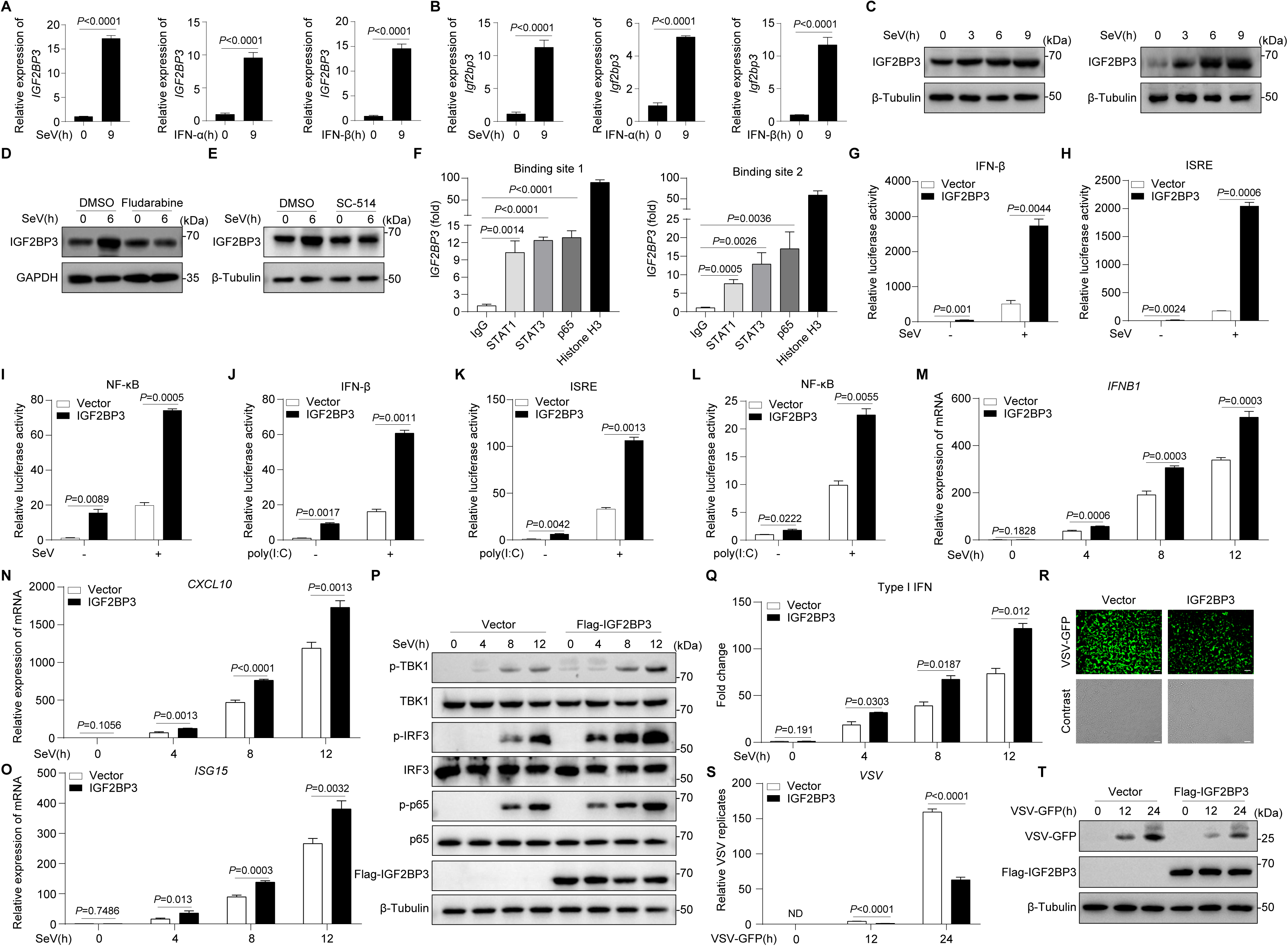
IGF2BP3 is a virus-inducible m^6^A reader that amplifies antiviral signaling. **A,B**, qRT-PCR of the relative *IGF2BP3* mRNA expression in THP-1 cells (A) or iBMDMs (B) stimulated with SeV, IFN-α, or IFN-β for 9 h. **C**, IB analysis of IGF2BP3 protein levels in THP-1 cells and iBMDMs infected with SeV. **D,E**, IB analysis of IGF2BP3 protein levels in THP-1 cells pretreated for 2 h with DMSO or fludarabine (D), or DMSO or SC-514 (E), followed by SeV infection and harvest at 0 and 6 h post-infection. **F**, ChIP–qPCR of STAT1, STAT3, or p65 occupancy at two IGF2BP3 promoter regions in HEK293T cells using IgG as a negative control and histone H3 as a positive control; enrichment is shown relative to IgG. **G-I**, Relative luciferase activities of the IFN-β (G), ISRE (H), or NF-κB (I) reporters in HEK293T cells transfected with empty vector or an IGF2BP3 expression plasmid, followed by SeV infection for 24 h. **J-L**, Relative luciferase activities of IFN-β (J), ISRE (K), or NF-κB (L) reporter in HEK293T cells transfected with empty vector or an IGF2BP3 expression plasmid, followed by poly(I:C) stimulation for 24 h. **M-O**, qRT-PCR of *IFNB1* (M), *CXCL10* (N), and *ISG15* (O) mRNA levels in HEK293T cells transfected with empty vector or an IGF2BP3 expression plasmid, at the indicated time points following SeV infection. **P**, IB analysis of phosphorylated and total TBK1, IRF3, and p65 in HEK293T cells transfected with empty vector or an IGF2BP3 expression plasmid, at the indicated time points following SeV infection. **Q**, Reporter-based analyses of bioactive IFN-I in supernatants from HEK293T cells transfected with empty vector or an IGF2BP3 expression plasmid and collected at the indicated times post-SeV infection, using 2fTGH-ISRE cells. **R-T**, VSV replication in HEK293T cells transfected with empty vector or an IGF2BP3 expression plasmid, followed by VSV-GFP infection, was assessed by GFP imaging at 24 h (R) and by qRT-PCR (VSV RNA; S) and IB (VSV-GFP; T) at 0, 12, and 24 h post-infection. Scale bars, 100 µm. Data are presented as mean ± SD. Statistical significance was evaluated using an unpaired two-tailed Student’s *t* test; exact *P* values are indicated in the panels. Data in C–E, P, and T are representative of three independent experiments with similar results.

To define the signaling pathways driving IGF2BP3 upregulation, we combined transcription factor binding-site prediction with pharmacological inhibition. PROMO(29) and JASPAR(30) analyses nominated STAT1, STAT3, and RELA (p65) as candidate regulators. In THP-1 cells, pharmacological inhibition of STAT1 (fludarabine) or IKKβ (SC-514) markedly reduced SeV-induced IGF2BP3 protein levels (Figure 1, D and E), indicating that both JAK–STAT and NF-κB signaling contribute to IGF2BP3 induction. Consistently, ChIP–qPCR revealed enrichment of STAT1, STAT3, and p65 at promoter regions flanking the transcription start site (TSS; −0.3 kb, binding site 1; and 0.8–1.1 kb, binding site 2) (Figure 1F). These data support direct promoter occupancy by STAT1, STAT3, and NF-κB p65 at the IGF2BP3 locus, providing a mechanistic basis for pathway-dependent induction.

Having established that IGF2BP3 is induced by antiviral signaling, we next investigated its functional role in innate immunity. IGF2BP3 overexpression significantly enhanced SeV- or poly(I:C)-induced activation of IFN-β, ISRE, and NF-κB luciferase reporters (Figure 1, G–L), with similar effects observed during HSV-1 infection (Supplemental Figure 1U). At the transcriptional level, IGF2BP3 overexpression increased SeV-induced *IFNB1*, *CXCL10*, and *ISG15* mRNA levels across multiple time points (Figure 1, M–O) and augmented phosphorylation of TBK1, IRF3, and p65 (Figure 1P), demonstrating potentiation of both the TBK1–IRF3 and NF-κB signaling axes. Moreover, supernatants from IGF2BP3-overexpressing cells contained increased bioactive IFN-I (Figure 1Q). Consistent with enhanced antiviral signaling, IGF2BP3 overexpression restricted vesicular stomatitis virus expressing green fluorescent protein (VSV-GFP) replication, as evidenced by reduced GFP fluorescence, lower viral RNA abundance, and decreased viral protein levels (Figure 1, R–T). Collectively, these results establish IGF2BP3 as a positive feedback regulator that is induced upon viral sensing and, in turn, potentiates antiviral signaling, augments IFN-β production, and restricts viral replication.

### IGF2BP3 loss-of-function attenuates antiviral innate immunity

To further define the physiological role of IGF2BP3 in antiviral innate immunity, we employed both knockdown and knockout approaches. Stable knockdown of *IGF2BP3* with two independent shRNAs in A549 cells (Supplemental Figure 2A) significantly diminished SeV-induced activation of IFN-β, ISRE, and NF-κB luciferase reporters (Supplemental Figure 2B). *IGF2BP3* knockdown also reduced SeV-induced *IFNB1* and *CXCL10* mRNA expression (Supplemental Figure 2, C and D), attenuated phosphorylation of TBK1, IRF3, and p65 (Supplemental Figure 2E), and consequently enhanced VSV-GFP replication (Supplemental Figure 2, F–H).

To establish a genetic loss-of-function model, we generated *Igf2bp3* KO iBMDMs and *IGF2BP3* KO HeLa cells using CRISPR–Cas9 (Figure 2A). Consistent with the knockdown phenotype, SeV infection induced markedly weaker activation of IFN-β, ISRE, and NF-κB reporters in *IGF2BP3* KO HeLa cells compared with wild-type (WT) controls (Figure 2B). In *Igf2bp3* KO iBMDMs, SeV-induced *Ifna4*, *Ifnb1*, and *Cxcl10* mRNA levels were significantly reduced across multiple time points (Figure 2, C–E), and phosphorylation of TBK1, IRF3, and p65 was diminished (Figure 2F), confirming impaired activation of both the TBK1–IRF3 and NF-κB pathways. Accordingly, *Igf2bp3* KO iBMDMs secreted less bioactive IFN-I (Figure 2G) and supported enhanced VSV-GFP replication, as evidenced by increased GFP fluorescence, higher viral RNA levels, and elevated viral protein abundance (Figure 2, H–J).

**Figure 2.**
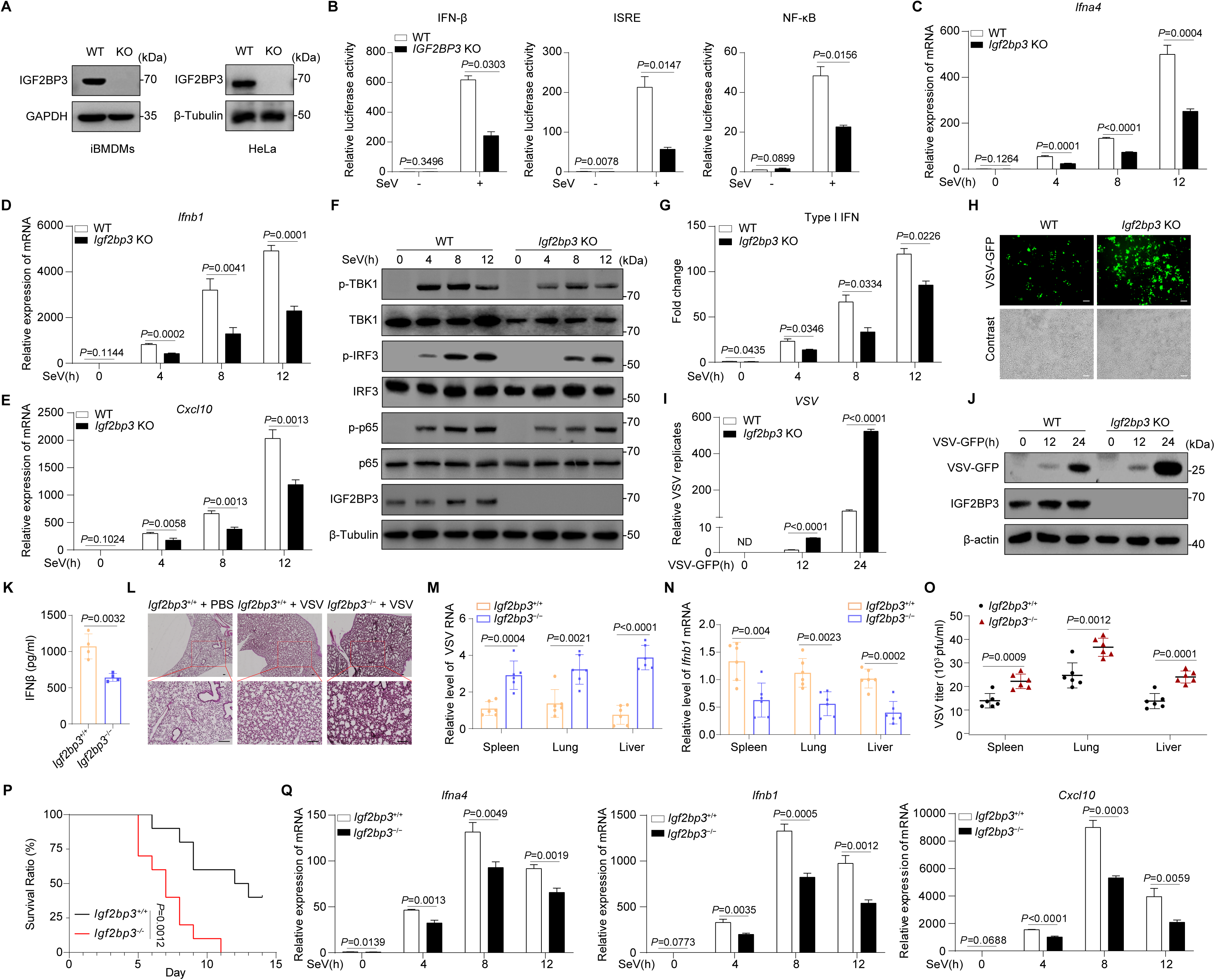
IGF2BP3 loss-of-function attenuates antiviral innate immunity. **A**, IB analysis validating *Igf2bp3* KO iBMDMs and *IGF2BP*3 KO HeLa cells. **B**, Relative luciferase activities of IFN-β, ISRE, and NF-κB reporters in WT and *IGF2BP3* KO HeLa cells infected with SeV for 24 h. **C-E**, qRT-PCR of *Ifna4* (C), *Ifnb1* (D), and *Cxcl10* (E) mRNA levels in WT and *Igf2bp3* KO iBMDMs at the indicated time points following SeV infection.. **F**, IB analysis of phosphorylated and total TBK1, IRF3, and p65 in WT and *Igf2bp3* KO iBMDMs at the indicated time points following SeV infection. **G**, Reporter-based analyses of bioactive IFN-I in supernatants from WT or *Igf2bp3* KO iBMDMs collected at the indicated times post–SeV infection, using L929-ISRE cells. **H-J**, VSV replication in WT and *Igf2bp3* KO iBMDMs infected with VSV-GFP was assessed by GFP imaging at 24 h (H) and by qRT-PCR (VSV RNA; I) and IB (VSV-GFP; J) at 0, 12, and 24 h post-infection. Scale bars, 100 µm. **K**, ELISA of serum IFN-β levels in *Igf2bp3*^+/+^ and *Igf2bp3*^−/−^ mice at 24 h post-VSV infection (n = 4 mice per group). **L**, Representative H&E-stained lung sections from *Igf2bp3*^+/+^ and *Igf2bp3*^−/−^ mice at 24 h post-VSV infection. Scale bars, 100 µm. **M,N**, qRT-PCR of VSV RNA abundance (M) and *Ifnb1* mRNA (N) in spleen, lung, and liver from *Igf2bp3*^+/+^ and *Igf2bp3*^−/−^ mice at 24 h post-VSV infection (n = 6 mice per group). **O**, Plaque assays quantifying infectious VSV titers in spleen, lung, and liver from *Igf2bp3*^+/+^ and *Igf2bp3*^−/−^ mice at 24 h post-VSV infection. **P**, Survival of *Igf2bp3^+/+^* mice and *Igf2bp3^−/−^*mice (n = 10 mice per group) at various times after tail-vein injection with VSV (1 × 10^8^ pfu/g). **Q**, qRT-PCR of *Ifna4, Ifnb1* and *Cxcl10* expression in peritoneal macrophages from *Igf2bp3*^+/+^ and *Igf2bp3*^−/−^ mice at the indicated time points following SeV infection.. Data are presented as mean ± SD. Statistical significance was determined by unpaired two-tailed Student’s *t* test; exact *P* values are indicated in the panels. Statistical significance in Figure 2Q was determined by the log-rank (Mantel–Cox) test. Data in A, F, and J are representative of three independent experiments with similar results.

To assess the physiological relevance of IGF2BP3 in antiviral defense *in vivo*, we established an acute VSV infection model using *Igf2bp3*^+/+^ and *Igf2bp3*^−/−^ mice (Supplemental Figure 2I). At 24 h post-infection, serum IFN-β levels were significantly lower in *Igf2bp3*^−/−^ mice (Figure 2K). Histological analysis revealed increased inflammatory infiltration in the lungs of *Igf2bp3*^−/−^ mice (Figure 2L). Consistently, qRT-PCR analysis of spleen, lung, and liver tissues showed elevated VSV RNA levels (Figure 2M) and reduced *Ifnb1* expression (Figure 2N) in *Igf2bp3*^−/−^ mice. Plaque assays confirmed higher viral titers in these organs (Figure 2O). Survival analysis further revealed increased susceptibility to viral infection in *Igf2bp3^−/−^* mice compared with WT controls (Figure 2P). Moreover, analysis of primary peritoneal macrophages following SeV infection showed that *Igf2bp3* deficiency markedly blunted the induction of *Ifna4*, *Ifnb1*, and *Cxcl10* expression at the indicated time points (Fig. 2R). Consistently, phosphorylation of TBK1, IRF3, and p65 was markedly reduced (Supplemental Fig. 2J), indicating impaired activation of both the TBK1–IRF3 and NF-κB pathways.

Collectively, these results demonstrate that endogenous IGF2BP3 is required for optimal antiviral signaling, IFN-I production, and restriction of RNA virus replication *in vitro* and *in vivo*.

### IGF2BP3 targets *MAVS* and *TBK1* transcripts to potentiate antiviral signaling

Given that IGF2BPs function as m^6^A readers(11, 31), we first tested whether METTL3-dependent m^6^A modification is required for IGF2BP3-mediated enhancement of antiviral signaling. The *METTL3* knockdown efficiency was confirmed (Supplemental Figure 3A).IGF2BP3 overexpression increased SeV-induced IFN-β reporter activity, whereas concomitant *METTL3* knockdown attenuated this enhancement (Figure 3A; Supplemental Figure 3C).

**Figure 3.**
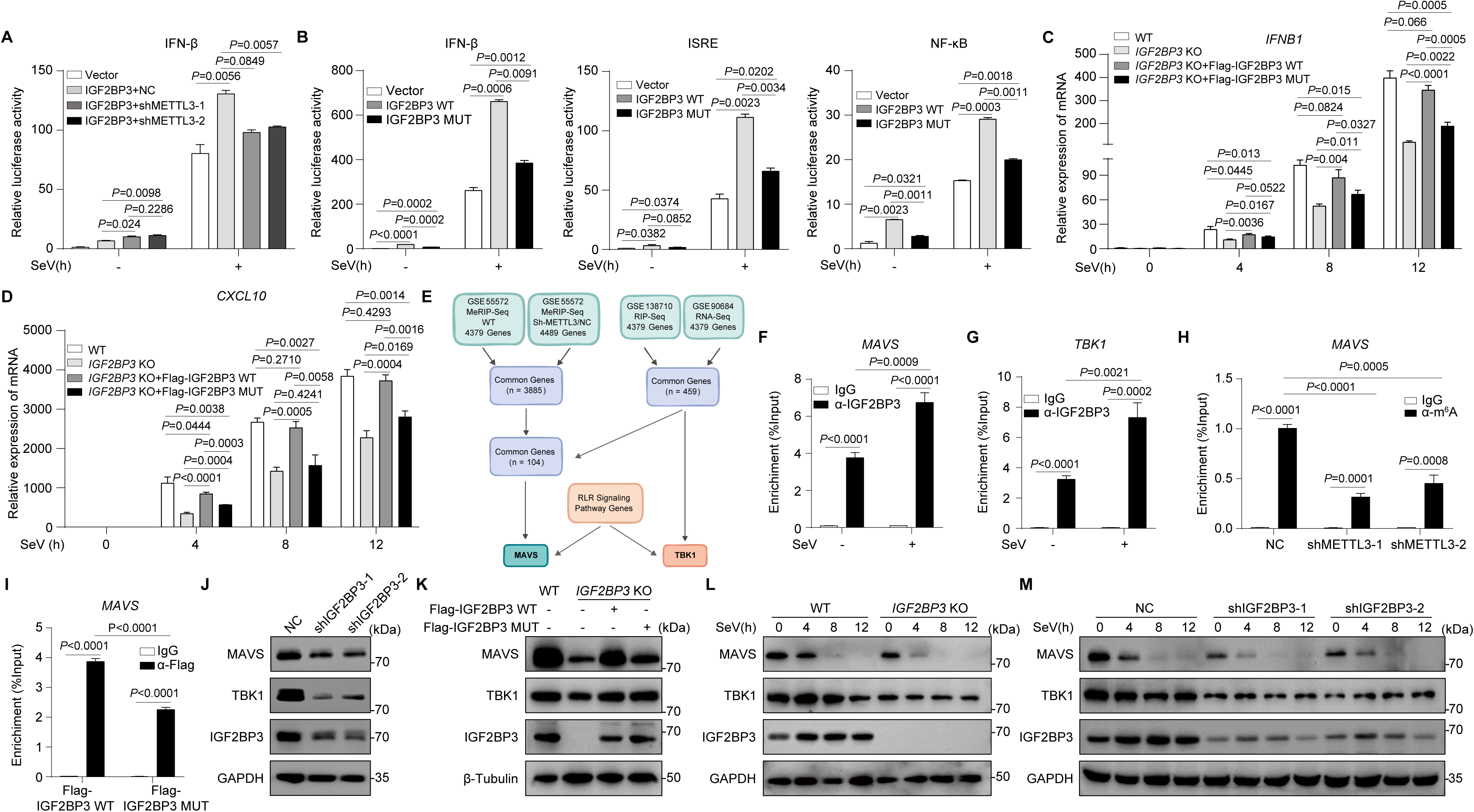
IGF2BP3 targets *MAVS* and *TBK1* transcripts to potentiate antiviral signaling. **A**, Relative luciferase activity of the IFN-β reporter in *IGF2BP3* KO HeLa cells transfected with empty vector or an IGF2BP3 expression plasmid together with control shRNA, shMETTL3-1, or shMETTL3-2, followed by SeV infection for 24 h. **B**, Relative luciferase activities of IFN-β, ISRE, and NF-κB reporters in *IGF2BP3* KO HeLa cells transfected with empty vector, IGF2BP3 WT, or an IGF2BP3 mutant, followed by SeV infection for 24 h. **C,D**, qRT-PCR of *IFNB1* (C) and *CXCL10* (D) mRNA levels in *IGF2BP3* KO HeLa cells reconstituted with empty vector, IGF2BP3 WT, or the IGF2BP3 mutant, infected with SeV and harvested at the indicated times post-infection. **E**, Integrative analyses of MeRIP-seq (GSE55572), RIP-seq (GSE138710), and RNA-seq (GSE90684) datasets identified *MAVS* and *TBK1* as candidate IGF2BP3 targets with relevance to the RLR signaling pathway. **F,G**, RIP–qRT-PCR of IGF2BP3-associated *MAVS* (F) and *TBK1* (G) transcripts in HEK293T cells with or without SeV infection using control IgG or anti-IGF2BP3 antibody. **H**, M^6^A-RIP–qRT-PCR of *MAVS* mRNA enrichment in HEK293T cells transfected with control shRNA or shMETTL3-1/shMETTL3-2 using IgG or anti-m^6^A antibody. **I**, RIP–qRT-PCR of *MAVS* mRNA enrichment in *IGF2BP3* KO HeLa cells reconstituted with Flag–IGF2BP3 WT or Flag–IGF2BP3 mutant using IgG or anti-Flag antibody. **J**, IB analysis of MAVS and TBK1 protein levels in A549 cells stably expressing control shRNA or shIGF2BP3-1/shIGF2BP3-2. **K**, IB analysis of MAVS and TBK1 protein levels in WT HeLa cells, *IGF2BP3* KO HeLa cells, and *IGF2BP3* KO cells reconstituted with Flag–IGF2BP3 WT or Flag–IGF2BP3 mutant. **L,M**, IB analysis of MAVS protein levels in WT and *IGF2BP3* KO HeLa cells (L) or in A549 cells stably expressing control shRNA or shIGF2BP3-1/shIGF2BP3-2 (M) infected with SeV and harvested at the indicated times post-infection. Data are presented as mean ± SD. Statistical significance was determined by unpaired two-tailed Student’s *t* test; exact *P* values are indicated in the panels. Data in J–M are representative of three independent experiments with similar results.

To directly assess the contribution of m^6^A recognition, we employed an IGF2BP3 mutant carrying GXXG-to-GEEG substitutions in the KH3/KH4 domains (Supplemental Figure 3B), which have been shown to disrupt m6A binding(11). Compared with WT IGF2BP3, the m^6^A-binding–defective mutant showed significantly reduced potentiation of SeV-induced IFN-β, ISRE, and NF-κB reporters, although reporter activity remained above vector controls (Figure 3B; Supplemental Figure 3D). Consistently, in *IGF2BP3* KO HeLa cells, reconstitution with WT IGF2BP3 restored SeV-induced *IFNB1* and *CXCL10* mRNA expression, whereas the m^6^A-binding mutant provided only partial rescue (Figure 3, C and D; Supplemental Figure 3E). These data indicate that m^6^A binding is required for the full antiviral activity of IGF2BP3 and IGF2BP3 promotes antiviral innate immune responses, at least in part, through its m^6^A-binding capacity.

To identify IGF2BP3 targets linked to RLR signaling, we integrated MeRIP-seq (GSE55572), IGF2BP3 RIP-seq (GSE138710), and RNA-seq (GSE90684) with RLR pathway gene set. This analysis highlighted *MAVS* as an m^6^A-enriched, IGF2BP3-associated transcript and *TBK1* as an IGF2BP3-associated candidate with relatively weak m^6^A enrichment (Figure 3E). RIP–qRT-PCR confirmed that IGF2BP3 associates with both *MAVS* and *TBK1* mRNAs, and that SeV infection strengthened these interactions (Figure 3, F and G; Supplemental Figure 3F). M^6^A-RIP–qRT-PCR further confirmed METTL3-dependent m^6^A enrichment on *MAVS* mRNA (Figure 3H), whereas *TBK1* exhibited only minimal m^6^A enrichment that was not significantly affected by *METTL3* knockdown (Supplemental Figure 3G). Accordingly, the m^6^A-binding–defective IGF2BP3 mutant exhibited reduced association with *MAVS* mRNA compared with WT IGF2BP3 (Figure 3I; Supplemental Figure 3I), while both forms associated with *TBK1* mRNA to a similar extent (Supplemental Figure 3H).

At the protein level, *IGF2BP3* knockdown in A549 cells reduced MAVS and TBK1 protein levels (Figure 3J). Similarly, *IGF2BP3* KO in HeLa cells decreased both proteins, and WT IGF2BP3 reconstitution restored their levels to near WT. In contrast, the m^6^A-binding defective mutant preferentially impaired restoration of MAVS, whereas TBK1 recovery was largely preserved (Figure 3K). Time-course analysis following SeV infection further revealed persistently reduced MAVS protein abundance in *IGF2BP3* KO cells (Figure 3L) and during early infection in *IGF2BP3* knockdown A549 cells (Figure 3M).

Collectively, these results support a model in which IGF2BP3 regulates *MAVS* predominantly in an m^6^A-dependent manner, whereas its regulation of *TBK1* is largely m^6^A-independent.

### IGF2BP3 stabilizes *MAVS* and *TBK1* mRNAs and facilitates their translation

Given that IGF2BP proteins are established regulators of mRNA stability(11, 31), we examined whether IGF2BP3 controls *MAVS* mRNA decay. Actinomycin D (ActD) chase experiments showed that *IGF2BP3* knockdown accelerated *MAVS* mRNA decay (Figure 4A) , *IGF2BP3* KO similarly reduced *MAVS* mRNA stability (Figure 4B), and IGF2BP3 overexpression prolonged *MAVS* mRNA persistence during ActD treatment (Figure 4C), indicating that IGF2BP3 stabilizes *MAVS* transcripts.

**Figure 4.**
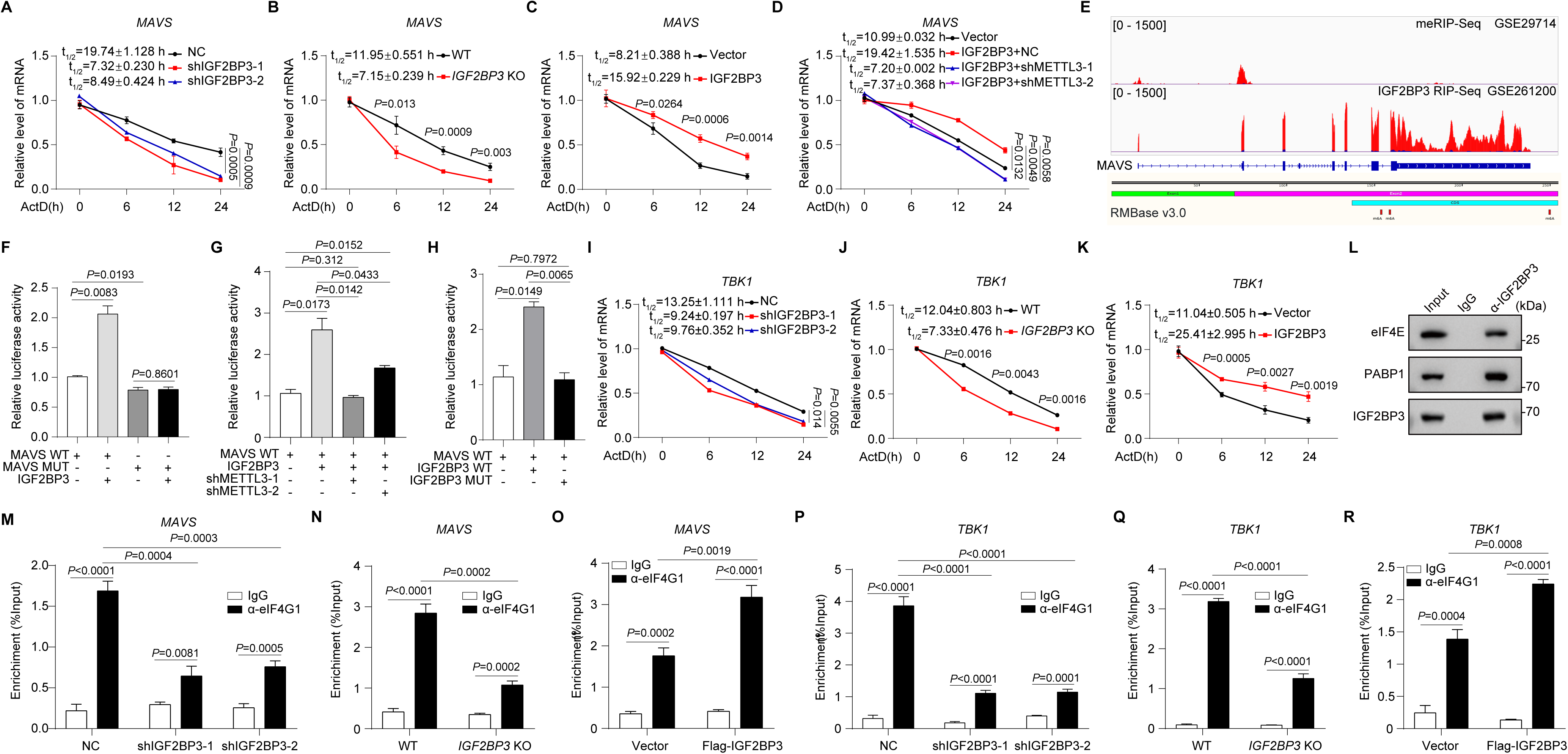
IGF2BP3 stabilizes *MAVS* and *TBK1* mRNAs and facilitates their translation. **A**-**D**, ActD-chase qRT-PCR of *MAVS* mRNA decay in A549 cells stably expressing shNC or shIGF2BP3-1/shIGF2BP3-2 (A), in WT and *IGF2BP3* KO HeLa cells (B), in HEK293T cells transfected with empty vector or IGF2BP3 (C), or co-transfected with IGF2BP3 plus control shRNA or shMETTL3-1/shMETTL3-2 (D), harvested at 0, 6, 12, and 24 h after ActD treatment (4 μg/mL). **E**, Schematic representation and IGV tracks showing m^6^A and IGF2BP3-binding peaks across the *MAVS* transcript, based on MeRIP-seq (GSE29714) and IGF2BP3 RIP-seq (GSE261200) datasets, with predicted residues annotated based on RMBase v3.0. **F-H**, Relative luciferase activities of pGL3- promoter-MAVS WT or m^6^A-site-mutant reporters in HEK293T cells with or without IGF2BP3 overexpression (F), with IGF2BP3 plus control shRNA or shMETTL3-1/shMETTL3-2 (G), or expressing IGF2BP3 WT versus an m^6^A-recognition-defective mutant (H). **I-K**, ActD-chase qRT-PCR of *TBK1* mRNA decay in A549 cells stably expressing shNC or shIGF2BP3-1/shIGF2BP3-2 (I), in WT and *IGF2BP3* KO HeLa cells (J), or in HEK293T cells transfected with empty vector or IGF2BP3 (K), harvested at 0, 6, 12, and 24 h after ActD treatment (4 μg/mL). **L**, Endogenous co-IP of IGF2BP3 association with eIF4E and PABP1 in HEK293T cells. Data are representative of three independent experiments with similar results. **M-R**, RIP–qRT-PCR of eIF4G1-associated *MAVS* mRNA in A549 cells stably expressing shNC or shIGF2BP3-1/shIGF2BP3-2 (M), in WT and *IGF2BP3* KO HeLa cells (N), and in HEK293T cells transfected with vector or IGF2BP3 (O), and eIF4G1-associated *TBK1* mRNA in the same settings (P–R). Data are presented as mean ± SD. Statistical significance was determined by unpaired two-tailed Student’s *t* test; exact *P* values are indicated in the panels.

Because IGF2BP3 association with *MAVS* mRNA depended partly on its m^6^A-binding capacity (Figure 3I), we next tested whether m^6^A is required for IGF2BP3-mediated *MAVS* stabilization. *METTL3* knockdown attenuated IGF2BP3-driven maintenance of *MAVS* mRNA during ActD chase, resulting in reduced *MAVS* mRNA levels at 6, 12, and 24 h (Figure 4D).

To identify potential m^6^A sites mediating IGF2BP3 binding of *MAVS* mRNA, we integrated MeRIP-seq (GSE29714) and IGF2BP3 RIP-seq (GSE261200) datasets. IGV (Integrative Genomics Viewer) analysis revealed prominent m^6^A enrichment on *MAVS* exon 2 and robust IGF2BP3-binding peaks across multiple exons, including exon 2; RMBase v3.0 predicted three putative m^6^A sites within exon 2 (Figure 4E).

To functionally validate whether IGF2BP3 regulates *MAVS* through these predicted m^6^A sites, we constructed luciferase reporters containing a 117-nt fragment of *MAVS* exon 2 in the 3′UTR (WT), or a mutant in which the predicted m^6^A-site adenosines were substituted with cytosine (Supplemental Figure 4A). IGF2BP3 overexpression significantly increased luciferase activity from the WT reporter but not from the mutant reporter (Figure 4F), indicating that IGF2BP3 regulation of this *MAVS* region requires intact m^6^A sites. This enhancement was attenuated by *METTL3* knockdown (Figure 4G) and abolished by the m^6^A-binding-defective IGF2BP3 mutant (Figure 4H), demonstrating that IGF2BP3 regulates *MAVS* mRNA through an m^6^A-dependent mechanism acting through these three sites in exon 2.

We next examined whether IGF2BP3 similarly regulates *TBK1* mRNA stability. We also analyzed the *TBK1* mRNA using the same dataset shown in Figure 4E. IGV analysis revealed few or no significant m^6^A peaks across the TBK1 transcript (Supplemental Figure 4B), consistent with our previous findings in Figure 3. ActD chase experiments demonstrated that *IGF2BP3* knockdown in A549 cells or *IGF2BP3* KO in HeLa cells reduced *TBK1* mRNA persistence, whereas IGF2BP3 overexpression prolonged *TBK1* mRNA stability (Figure 4, I–K), indicating that IGF2BP3 also stabilizes *TBK1* transcripts.

Prior work has suggested that IGF2BP family proteins can enhance translation of target mRNAs(11), but the underlying mechanism remains unclear. An independent proteomic study indicated that IGF2BP3 likely associates with components of the eIF4F/PABP-associated initiation complex, including eIF4E and PABP1(32). We therefore hypothesized that IGF2BP3 promotes translation by interacting with initiation factors and increasing the occupancy of its target mRNAs within the eIF4F/PABP-associated initiation complex, thereby promoting their translation. To test this hypothesis, we performed co-immunoprecipitation (co-IP) and found that IGF2BP3 associated with the translation initiation factors eIF4E and with PABP1 (Figure 4L), but not detectably with eIF4G1 (Supplemental Figure 4C).

Using an established eIF4G1-RIP strategy(33–35), we found that *IGF2BP3* knockdown reduced eIF4G1-associated *MAVS* mRNA in A549 cells (Figure 4M) without affecting *GAPDH* (Supplemental Figure 4D). Similar reductions were observed in *IGF2BP3* KO HeLa cells (Figure 4N), whereas IGF2BP3 overexpression increased eIF4G1-associated *MAVS* mRNA in HEK293T cells (Figure 4O) without affecting *GAPDH* (Supplemental Figure 4, E and F). Parallel analyses demonstrated that IGF2BP3 similarly modulates eIF4G1-associated *TBK1* mRNA (Figure 4, P–R).

Collectively, these findings demonstrate that IGF2BP3 stabilizes both *MAVS* and *TBK1* mRNAs and promotes their association with the eIF4G1-containing translation initiation machinery, thereby facilitating their translation.

### IGF2BP3 associates with stress granule components and promotes RIG-I recognition of viral RNA

Stress granules (SGs) are cytoplasmic condensates that form in response to diverse cellular stresses, including viral infection(36). Although IGF2BP family members have been reported to localize to SGs induced by osmotic stress (37) or heat shock(11), their role in avSGs remains unclear. We found that upon VSV infection or poly(I:C) stimulation, IGF2BP3 redistributed from a diffuse cytoplasmic pattern to punctate foci that co-localized with the canonical SG marker G3BP1 (Figure 5A). Co-IP assays showed that IGF2BP3 interacted with G3BP1, and this interaction was enhanced upon VSV infection (Figure 5B). *In vitro* binding assays further supported a direct IGF2BP3–G3BP1 interaction (Figure 5C). IGF2BP3 also associated with the RNA sensor RIG-I, with increased association following VSV infection (Figure 5D).

**Figure 5.**
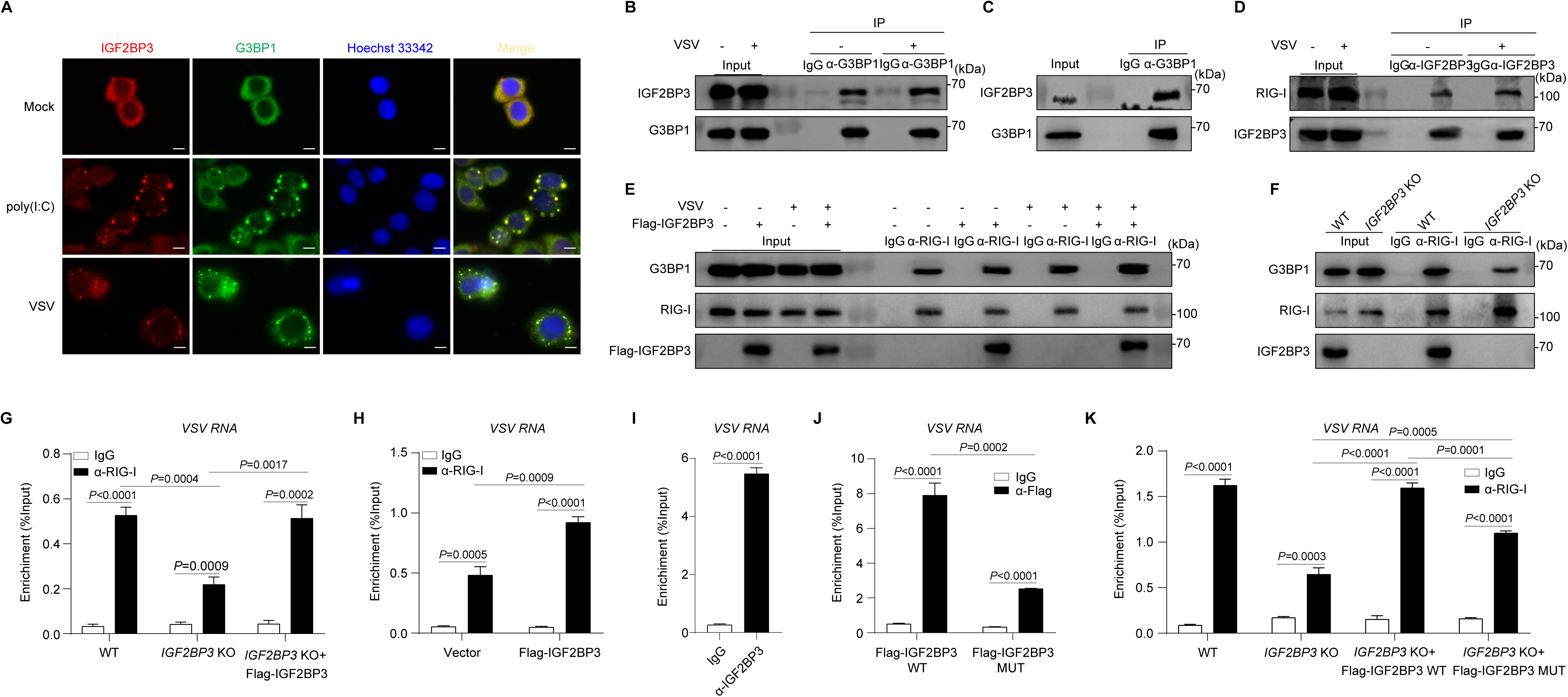
IGF2BP3 associates with stress granule components and promotes RIG-I recognition of viral RNA. **A**, Immunofluorescence of IGF2BP3 and G3BP1 localization in HeLa cells left untreated or stimulated with VSV or poly(I:C) for 6 h. Scale bars, 10 µm. **B**, Endogenous co-IP of IGF2BP3 association with G3BP1 in HEK293T cells mock-treated or infected with VSV for 6 h. **C**, *In vitro* binding assays assessing the IGF2BP3–G3BP1 interaction by IB using proteins synthesized by the TnT® Quick Coupled Transcription/Translation System and immunoprecipitated with an anti-G3BP1 antibody. **D**, Endogenous co-IP of RIG-I association with IGF2BP3 in HEK293T cells mock-treated or infected with VSV for 6 h. **E,F**, Co-IP of the RIG-I–G3BP1 interaction in HEK293T cells transfected with empty vector or Flag–IGF2BP3 (E) and in WT versus *IGF2BP3* KO HeLa cells (F) following mock treatment or VSV infection for 6 h. **G-K**, RIP–qRT-PCR of VSV RNA associated with RIG-I in WT, *IGF2BP3* KO, and reconstituted *IGF2BP3* KO HeLa cells (G) and in HEK293T cells transfected with empty vector or Flag–IGF2BP3 (H), and of VSV RNA associated with IGF2BP3 in HEK293T cells (I), as well as Flag-RIP–qRT-PCR in *IGF2BP3* KO HeLa cells reconstituted with Flag–IGF2BP3 WT or mutant (J) and RIG-I RIP–qRT-PCR analyses in WT/KO/reconstituted HeLa cells (K). Data are presented as mean ± SD. Statistical significance was determined by unpaired two-tailed Student’s *t* test; exact *P* values are indicated in the panels. Data in B–F are representative of three independent experiments with similar results.

Since avSGs serve as platforms for RIG-I-mediated viral RNA sensing and downstream signaling(25, 38), we asked whether IGF2BP3 modulates RIG-I–G3BP1 complex formation. IGF2BP3 overexpression increased the RIG-I–G3BP1 association both at baseline and after VSV infection (Figure 5E), whereas *IGF2BP3* KO reduced this interaction (Figure 5F), suggesting that IGF2BP3 facilitates assembly of the RIG-I–G3BP1 complex. To determine whether IGF2BP3 influences RIG-I binding to viral RNA, we performed RIP assays. RIG-I-associated VSV RNA was reduced in *IGF2BP3* KO cells and restored by IGF2BP3 reconstitution (Figure 5G), whereas IGF2BP3 overexpression increased RIG-I-associated VSV RNA in HEK293T cells (Figure 5H). No enrichment of the negative control transcript *GAPDH* was detected (Supplemental Figure 4, G-I).

Anti-IGF2BP3 RIP further revealed enrichment of VSV RNA (Figure 5I). Given prior evidence that VSV RNA is m^6^A-modified(6, 39), we tested whether IGF2BP3 binding to VSV RNA requires m^6^A recognition. In *IGF2BP3* KO HeLa cells reconstituted with Flag-tagged constructs, both WT and m^6^A-binding–defective IGF2BP3 enriched VSV RNA, but the mutant showed significantly reduced association (Figure 5J; Supplemental Figure 4J), indicating a partial requirement for m^6^A recognition for maximal viral RNA binding. Consistently, RIG-I RIP showed reduced VSV RNA association in cells reconstituted with the m^6^A-binding mutant relative to WT reconstitution (Figure 5K; Supplemental Figure 4, K and L).

Collectively, these data demonstrate that IGF2BP3 is recruited to avSGs, directly interacts with G3BP1, associates with RIG-I, promotes RIG-I–G3BP1 complex formation, and enhances RIG-I binding to viral RNA, at least in part through its m^6^A-binding capacity.

### Curcumol targets IGF2BP3 to impair *MAVS* and *TBK1* mRNA stability and translation

Dysregulated IFN-I signatures are closely associated with SLE(40, 41). Given that IFN-I upregulates IGF2BP3 and that IGF2BP3 amplifies IFN-I production, we asked whether aberrant IGF2BP3 feedback might contribute to interferon-associated autoimmune inflammation. In two public SLE cohorts (GSE61635 and GSE49454), *IGF2BP3* expression was significantly elevated in patients relative to healthy controls, whereas *IGF2BP1* remained unchanged and *IGF2BP2* was reduced (Figure 6, A and B; Supplemental Figure 5, A–D). *IGF2BP3* expression positively correlated with IFN-I–related genes, and IGF2BP3-associated gene signatures were enriched for IFN-I and antiviral innate immune pathways (Supplemental Figure 5, E and F), suggesting a potential role for IGF2BP3 in sustaining the interferon-driven inflammatory program in SLE. We next evaluated IGF2BP3 in an interferonopathy-like autoimmune model. Immunoblotting showed that IGF2BP3 protein levels were increased in peritoneal macrophages from *Trex1* KO mice compared with WT controls (Figure 6C). The data suggested that IGF2BP3 may serve as a potential therapeutic target for interferon-associated autoimmune inflammation.

**Figure 6.**
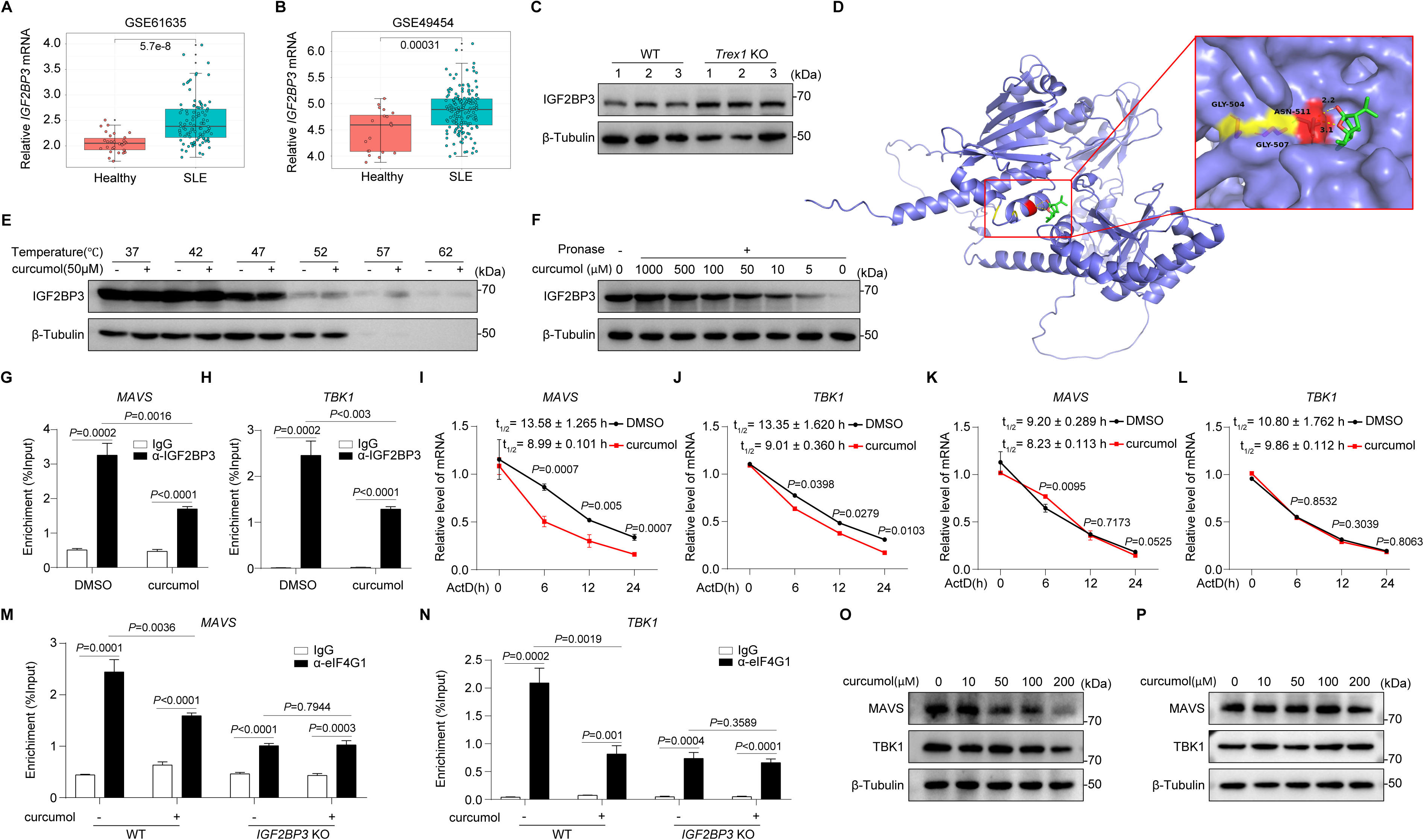
Curcumol targets IGF2BP3 to impair *MAVS* and *TBK1* mRNA stability and translation. **A,B**, *IGF2BP3* expression levels in healthy controls and SLE patients from public datasets GSE61635 (A) and GSE49454 (B). **C**, IB analysis of IGF2BP3 protein levels in peritoneal macrophages isolated from WT and *Trex1* KO mice. **D**, Molecular docking analyses predicting the curcumol-IGF2BP3 interaction using AutoDock4, with docking poses rendered in PyMOL. **E**, DARTS analyses of IGF2BP3 protease resistance in HEK293T lysates pre-incubated with DMSO or curcumol (5–1000 μM) for 1 h and digested with pronase (5 μg/mL, 1 h); β-tubulin served as a loading control. **F**, CETSA analyses of IGF2BP3 thermal stability in HEK293T cells treated with DMSO or curcumol (50 μM) for 24 h and heated from 37°C to 62°C; β-tubulin served as a control. **G,H**, RIP–qRT-PCR of IGF2BP3-associated *MAVS* (G) and *TBK1* (H) transcripts in HEK293T cells treated with DMSO or curcumol (50 μM) for 24 h using IgG or anti-IGF2BP3 antibody. **I,J**, ActD-chase qRT-PCR of *MAVS* (I) and *TBK1* (J) mRNA decay in HeLa cells pretreated with DMSO or curcumol (50 μM) for 24 h and harvested at 0, 6, 12, and 24 h after ActD treatment (4 μg/mL). **K,L**, ActD-chase qRT-PCR of *MAVS* (K) and *TBK1* (L) mRNA decay in *IGF2BP3* KO HeLa cells pretreated with DMSO or curcumol (50 μM) for 24 h and harvested at 0, 6, 12, and 24 h after ActD treatment (4 μg/mL). **M,N**, RIP–qRT-PCR of eIF4G1-associated *MAVS* (M) and *TBK1* (N) mRNAs in WT and *IGF2BP3* KO HeLa cells treated with DMSO or curcumol (50 μM) for 24 h. **O,P**, IB analysis of MAVS and TBK1 protein levels in WT HeLa cells (O) or *IGF2BP3* KO HeLa cells (P) treated with curcumol (0–200 μM) for 24 h. Data are presented as mean ± SD. Statistical significance was determined by unpaired two-tailed Student’s *t* test; exact *P* values are indicated in the panels. Data in C, E, F, O, and P are representative of three independent experiments with similar results.

Curcumol has been reported to reduce IGF2BP3 protein levels in cancer contexts(42), but direct evidence of its IGF2BP3 targeting has been limited. Given that IGF2BP3 is a potential therapeutic target, we explored whether curcumol could pharmacologically modulate IGF2BP3 activity. Molecular docking predicted favorable binding of curcumol to IGF2BP3 (ΔG = −5.05 kcal/mol; Figure 6D). Experimentally, DARTS assays showed that curcumol increased the protease resistance of IGF2BP3 in a dose-dependent manner without affecting β-tubulin (Figure 6E). CETSA further demonstrated that curcumol enhanced the thermal stability of IGF2BP3 (Figure 6F), confirming IGF2BP3 target engagement in cells.

Functionally, curcumol reduced IGF2BP3–mRNA interactions, as RIP–qRT-PCR revealed decreased association of IGF2BP3 with *MAVS* and *TBK1* mRNAs (Figure 6, G and H) and with the known target *MYC* (Supplemental Figure 5, G and H). Curcumol also destabilized *MAVS* and *TBK1* transcripts during ActD chase in WT HeLa cells (Figure 6, I and J), and similarly reduced *MYC* mRNA stability (Supplemental Figure 5I); however, these effects were largely lost in *IGF2BP3* KO cells (Figure 6, K and L; Supplemental Figure 5J). Moreover, curcumol decreased eIF4G1-associated *MAVS*, *TBK1*, and *MYC* mRNAs in WT cells but not in *IGF2BP3* KO cells (Figure 6, M and N; Supplemental Figure 5, K and L), indicating that its effects on destabilizing IGF2BP3-target transcripts and inhibiting translation complex recruitment were IGF2BP3-dependent. Consistently, curcumol dose-dependently reduced MAVS and TBK1 protein levels in WT cells (Figure 6O) but had minimal effects in *IGF2BP3* KO cells (Figure 6P).

Collectively, these data demonstrate that curcumol directly engages IGF2BP3 and suppresses its binding to *MAVS* and *TBK1* mRNAs, thereby impairing transcript stability and translation-complex recruitment and reducing MAVS and TBK1 protein expression.

### Curcumol ameliorates interferon-driven autoimmune pathology

The above findings indicate that IGF2BP3 is a potential therapeutic target for interferon-associated autoimmune inflammation (Figure 6, A–C). We therefore assessed the effects of curcumol-mediated IGF2BP3 inhibition *in vitro* and *in vivo*. In *Trex1* KO peritoneal macrophages, curcumol treatment significantly reduced *Ifnb1*, *Il6,* and *Isg15* expression compared with DMSO (Figure 7A), indicating suppression of IFN-I and inflammatory gene expression *in vitro*. We next treated *Trex1* KO mice with curcumol daily for 2 weeks (Figure 7B). Curcumol treatment significantly prolonged survival (Figure 7C), reduced splenomegaly and spleen index (Figure 7D), and decreased *Ifnb1*, *Il6*, and *Isg15* expression across multiple organs, including heart, liver, kidney, muscle, stomach, and tongue (Figure 7, E–J). Histological analyses further showed that curcumol attenuated inflammatory infiltration and tissue damage in these organs (Figure 7K).

**Figure 7.**
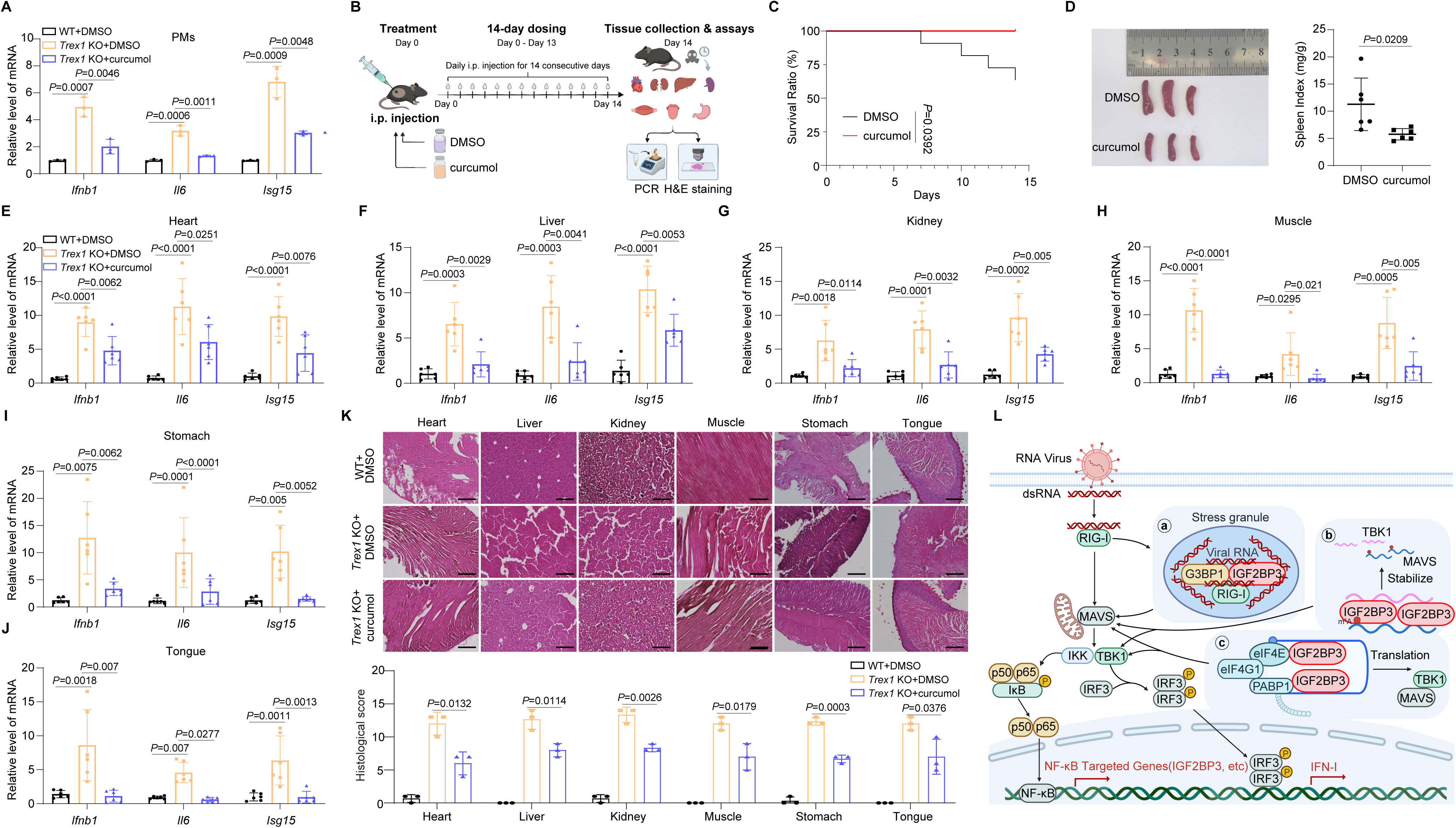
Curcumol ameliorates interferon-driven autoimmune pathology. **A**, qRT-PCR of *Ifnb1*, *Il6*, and *Isg15* mRNA expression in peritoneal macrophages from WT mice (DMSO) and *Trex1* KO mice treated with DMSO or curcumol (50 μM) for 24 h. **B**, Schematic of the curcumol treatment in *Trex1* KO mice (daily intraperitoneal injection of DMSO or curcumol (50 μg/g) for 14 days; tissues collected 24 h after the final dose). **C**, Survival of *Trex1* KO mice treated with DMSO or curcumol (50 μg/g daily for 2 weeks; n = 10 per group), with randomization and investigator blinding as indicated; Kaplan–Meier curves were compared by log-rank test. **D**, Representative spleen images and spleen index (n = 6 mice per group) quantification of *Trex1* KO mice treated with DMSO or curcumol. **E-J**, qRT-PCR of *Ifnb1*, *Il6*, and *Isg15* mRNA levels in heart (E), liver (F), kidney (G), muscle (H), stomach (I), and tongue (J) from *Trex1* KO mice treated with DMSO or curcumol (n = 6 mice per group). **K**, Histological analyses including representative H&E staining (upper panel) and blinded pathology scoring (bottom panel) of heart, liver, kidney, muscle, stomach, and tongue from WT mice and DMSO- or curcumol-treated *Trex1* KO mice. Scale bars, 100 µm. **L**, Working model for IGF2BP3-mediated regulation of antiviral innate immunity. IGF2BP3 binds and stabilizes *MAVS* and *TBK1* mRNAs and facilitates their recruitment to the eIF4F/PABP-associated initiation complex, thereby enhancing translation. Upon RNA virus infection, IGF2BP3 relocalizes to avSGs, where it interacts with G3BP1 and RIG-I and promotes assembly of the RIG-I–G3BP1 complex to facilitate viral RNA sensing. IGF2BP3 amplifies virus-induced activation of the TBK1–IRF3 and NF-κB pathways, promoting IFN-I production, which in turn induces *IGF2BP3* expression to establish a feedforward loop. Data are presented as mean ± SD. Statistical significance was determined by unpaired two-tailed Student’s *t* test for pairwise comparisons and by log-rank test for survival; exact *P* values are indicated in the panels.

Collectively, these data demonstrate that pharmacological inhibition of IGF2BP3 by curcumol ameliorates IFN-I-driven pathology and improves disease outcomes in *Trex1* KO mice.

## Discussion

The RNA-binding protein IGF2BP3 has been widely recognized as a key m^6^A reader involved in cancer progression and stem cell maintenance(11, 17). However, its role in antiviral innate immunity has remained largely unexplored. Our study establishes IGF2BP3 as a critical positive regulator of antiviral signaling, functioning through multifaceted mechanisms that integrate RNA metabolism, translational control, and stress granule–mediated viral RNA sensing (Figure 7L).

Previous studies have identified several m^6^A readers that regulate innate immunity, including YTHDF proteins that modulate the stability of interferon-stimulated genes(7, 43, 44). However, direct regulation of *MAVS* and *TBK1* by specific m^6^A readers has not been systematically characterized. Our work demonstrates that IGF2BP3 directly targets *MAVS* and *TBK1* mRNAs, with distinct m^6^A dependency: *MAVS* regulation requires intact m^6^A binding, while *TBK1* regulation appears largely m^6^A-independent. This differential mechanism suggests that IGF2BP3 employs distinct RNA-binding modes to coordinate the expression of key signaling components in the antiviral pathway.

Beyond *MAVS* and *TBK1*, IGF2BP3 has been reported to regulate numerous transcripts involved in cell proliferation and metabolism, including *MYC*, *IGF2*, and *HMGA2*(45–47). Our study expands this repertoire to include critical antiviral signaling molecules, positioning IGF2BP3 as a broad regulator that links cellular growth control with immune defense.

Recent studies have highlighted the importance of SGs as platforms for antiviral sensing, with proteins such as G3BP1 facilitating RIG-I activation (38, 48, 49). Our findings reveal that IGF2BP3 is recruited to virus-induced SGs, interacts with both G3BP1 and RIG-I, and promotes RIG-I–G3BP1 complex formation, thereby enhancing RIG-I binding to viral RNA. Published studies have established that VSV RNA carries m^6^A modifications, supported by transcriptome-wide mapping and functional validation (6, 39). Our findings add a novel layer of regulation to stress granule–mediated antiviral defense. Importantly, our findings indicate that IGF2BP3 binds VSV RNA in an m^6^A-dependent manner, providing a molecular basis for coupling m^6^A readout to viral RNA recognition. Moreover, prior work suggests that m^6^A modification is one determinant of self versus non-self RNA discrimination, and that m^6^A marks on viral RNA can dampen RIG-I sensing (22, 50, 51). In this context, the ability of IGF2BP3 to read m^6^A on viral RNA while facilitating RIG-I engagement may represent a host strategy to enhance antiviral defense.

A significant finding of our study is the identification of a feedforward loop in which viral infection or interferon stimulation upregulates IGF2BP3, which in turn amplifies antiviral responses. While this circuit provides robust defense against pathogens, its dysregulation may contribute to autoimmune pathology. Excessive IFN-I production is a hallmark of systemic autoimmune diseases such as SLE(41, 52, 53). Our observation that IGF2BP3 is upregulated in SLE patients and *Trex1* KO mice suggests it may participate in sustaining pathological interferon signatures. Moreover, IGF2BP3 may lower the threshold for RIG-I recognition of self RNA, thereby promoting the initiation or exacerbation of autoimmunity. This connection motivated our investigation of pharmacological inhibition of IGF2BP3 by curcumol in interferon-driven pathology. I*n vivo*, curcumol treatment significantly ameliorated interferon-driven autoimmune pathologies.

The upregulation of IGF2BP3 in SLE patients and *Trex1* KO mice, along with the therapeutic efficacy of curcumol in suppressing interferonopathy, suggests that targeting IGF2BP3 may represent a promising strategy for treating interferon-associated autoimmune diseases. Current therapies for such conditions often involve broad immunosuppression (54), whereas targeting specific regulators like IGF2BP3 could offer more precise intervention with potentially fewer side effects. Our demonstration that curcumol engages IGF2BP3 and disrupts its function provides a pharmacological foundation for developing more selective IGF2BP3 inhibitors.

While our study provides substantial evidence for IGF2BP3’s role in antiviral immunity, several questions remain. The precise structural basis for the differential m^6^A dependency of IGF2BP3 toward *MAVS* versus *TBK1* mRNAs warrants further investigation. Moreover, the subcellular site at which IGF2BP3 stabilizes *MAVS* and *TBK1* mRNAs and promotes their translation during viral infection remains to be defined, whether this occurs within avSGs, other membrane-less compartments, or more broadly in the cytosol. The potential role of IGF2BP3 in modulating the translation efficiency of target mRNAs, either through interactions with additional translation-related factors or by participating in other stages of the translation process, also requires further exploration. Additionally, the coordination between IGF2BP3 and other m^6^A readers in antiviral contexts, as well as the possible variation in its functions across distinct cell types or viral infections, represents important avenues for future investigation. Existing literature has reported m^6^A modifications on *IFNB1* and *ISG15* in antiviral innate immune pathways (55–57). However, whether IGF2BP3 regulates other m^6^A-modified mRNAs that were not identified in our multi-omics screening remains an unresolved question, warranting further investigation. Growing evidence indicates that viral infections, particularly SARS-CoV-2 infection, are associated with an increased risk of new-onset autoimmune diseases(58), highlighting the pathological consequences of sustained innate immune activation and dysregulated type I interferon responses. Together with our findings that IGF2BP3 is upregulated in systemic lupus erythematosus and *Trex1* KO mice and promotes IFN-I–driven inflammation, these observations raise the possibility that aberrant IGF2BP3 activation may contribute to post-viral autoimmune manifestations, a possibility that merits further investigation.

In summary, our work identifies IGF2BP3 as a central amplifier of antiviral innate immunity that operates through coordinated regulation of RNA stability, translation efficiency, and stress granule–mediated viral RNA detection (Figure 7L). The feedforward loop involving virus/IFN-induced *IGF2BP3* expression and subsequent immune amplification represents a sophisticated regulatory mechanism that ensures effective antiviral defense while potentially contributing to autoimmune pathology when dysregulated. These findings not only advance our understanding of post-transcriptional regulation in innate immunity but also highlight IGF2BP3 as a potential therapeutic target for managing both viral infections and interferon-related autoimmune conditions.

## Materials and Methods

### Antibodies

Primary antibodies used in this study were: GAPDH (mouse mAb; Abmart, M20006S), β-actin (mouse mAb; Proteintech, 66009-1-Ig), β-tubulin (mouse mAb; Abmart, M30109S), Flag tag (mouse mAb; MBL, M185-3L), HA tag (mouse mAb; Abmart, M20003S), GFP (mouse pAb; MBL, MBL598), IGF2BP3 (rabbit pAb; Proteintech, 14642-1-AP), IGF2BP3 (mouse mAb; Santa Cruz, sc-390639), m^6^A (mouse mAb; Proteintech, 68055-1-Ig), NF-κB p65 (rabbit mAb; Cell Signaling Technology, 8242S), phospho-NF-κB p65 (Ser536) (rabbit mAb; Cell Signaling Technology, 3033S), G3BP1 (rabbit pAb; Proteintech, 13057-2-AP), eIF4E (rabbit mAb; STARTER, S0B6141), PABP1 (rabbit pAb; STARTER, S0B1354), TBK1 (rabbit pAb; Proteintech, 28397-1-AP), phospho-TBK1 (Ser172) (rabbit mAb; Cell Signaling Technology, 5483S), IRF3 (rabbit pAb; Proteintech, 11312-1-AP), phospho-IRF3 (Ser396) (rabbit pAb; Cell Signaling Technology, 4947S), Histone H3 (mouse mAb; Beyotime, Cat#AF0009), STAT3 mAb (ABclonal, A19566), STAT1 mAb (Cell Signaling Technology, 14995S), mouse IgG (Sigma-Aldrich, I5381), rabbit IgG (Sigma-Aldrich, I5006).

### Chemicals and reagents

curcumol (Yuanye, B20342), actinomycin D (Act D, Selleck, S8964), fludarabine (Selleck, S1491), SC-514 (Selleck, S4907), poly(I:C) (InvivoGen, tlrl-pic), recombinant human IFN-α (Novoprotein, C005), recombinant human IFN-β (Sino Biological, 10704-H02H), proteinase K (Biosharp, CP9191), Hoechst 33258 (Beyotime, C1011), 4% paraformaldehyde (PFA; Biosharp, BL539A), DMSO (Solarbio, D8371), BSA (Yeasen, 36101ES60).

### Cell culture

HEK293T, HeLa, A549, iBMDMs, and Vero cells were maintained in DMEM (Thermo Fisher Scientific, C11995500BT) supplemented with 10% heat-inactivated fetal bovine serum (FBS; CellMax, FSP500) and 1% penicillin-streptomycin (Solarbio, P1400). THP-1 cells and primary human peritoneal macrophages were cultured in RPMI-1640 (Thermo Fisher Scientific, C11875500BT) medium containing 10% FBS and antibiotics. Adherent cells were detached using 0.25% trypsin-EDTA (Macgene, CC017). All cells were incubated at 37 °C under 5% CO2.

### plasmids, and transfection

Flag-IGF2BP3-WT, Flag-IGF2BP3-MUT (carrying GXXG-to-GEEG substitutions in the KH3/KH4 domains), pGL3-Promoter–MAVS-WT and pGL3-Promoter–MAVS-MUT (the 17th, 22nd, and 113th adenosines were mutated to guanosines) were generated in this study using standard molecular cloning procedures. All constructs were verified by sanger sequencing prior to use. Plasmid DNA and short hairpin RNA (shRNA) transfections were performed using Lipo6000 (Beyotime, C0526) or polyethylenimine (PEI; Polysciences, 49553-93-7) according to the manufacturer’s instructions.

### Generation of knockdown and knockout cell lines

Stable knockdown cell lines were generated by lentiviral delivery of shRNAs. Lentiviral particles were produced in HEK293T cells by co-transfecting pLKO-based shRNA plasmids (pLKO.1 puro; Addgene #8453) with the packaging plasmid psPAX2 (Addgene #12260) and the envelope plasmid pMD2.G (Addgene #12259) using standard procedures. Viral supernatants were collected, clarified by filtration, and used to transduce target cells in the presence of polybrene (Beyotime, C0351), followed by puromycin selection (Solarbio, P8230) to obtain stable knockdown populations. Knockdown efficiency was validated by immunoblotting. The shRNA sequences targeting human IGF2BP3 were: shRNA1, 5′-GAAACTTCAGATACGAAATAT-3′; shRNA2, 5′-AATCGATGTCCACCGTAAAGA-3′.

Knockout cell lines were generated using a lentivirus-packaged CRISPR–Cas9 system. sgRNAs targeting the gene of interest were cloned into lentiCRISPRv2 (Addgene #52961), and lentiviruses were produced in HEK293T cells using psPAX2 (Addgene #12260) and pMD2.G (Addgene #12259) and subsequently used to transduce target cells in the presence of polybrene (Beyotime, C0351). After puromycin selection (Solarbio, P8230), single-cell clones were isolated by limiting dilution and expanded. Gene disruption was confirmed by immunoblotting and, where indicated, by genomic PCR followed by Sanger sequencing. The sgRNA target sequences were: human IGF2BP3 sgRNA, 5′-CTTACCTCCCACTGTAAATG-3′; and mouse Igf2bp3 sgRNA, 5′-ACTCGGTCCCTAAACGGCAG-3′.

### Virus amplification

SeV was propagated in 9-day-old SPF embryonated chicken eggs. After inoculation into the allantoic cavity and incubation at 37 °C for 72 h, allantoic fluid was harvested, clarified by centrifugation, aliquoted, and stored at −80 °C. VSV, VSV-GFP and HSV-1 were amplified in Vero cells. Upon extensive cytopathic effect, supernatant was collected, debris-removed by centrifugation at 3000 g for 10 min at 4 °C, and virus concentrated by ultracentrifugation at 71000 g for 1 h at 4 °C. Viral pellets were resuspended in Tris-HCl (pH 7.5, 10 mM) and stored at −80 °C.

### Plaque assay

Vero cells in 24-well plates were infected with serially diluted virus for 1 h, overlaid with 0.5% methylcellulose (Sigma-Aldrich, M0512), and incubated for 2–3 days. Cells were fixed with 4% PFA, stained with crystal violet (MREDA, M1415), and plaques were counted. Viral titers were calculated based on plaque counts and dilution factors.

### Dual-luciferase reporter assay

Cells in 24-well plates were transfected with firefly luciferase reporter plasmids (IFN-β, ISRE, or NF-κB) and pRL-SV40 (Renilla control) using PEI. After treatment, cells were lysed and luminescence was measured sequentially using Dual-Luc Pro Luciferase Reporter Assay Kit (TransDetect, FR203-01). All reporter assays were performed in duplicate andrepeated independently at least three times. Relative activity was calculated as the firefly/Renilla ratio and normalized to controls.

### Quantitative real-time PCR (qRT-PCR)

Total RNA was extracted using TRIzol reagent (APPLYGEN, R1030) according to the manufacturer’s instructions. Reverse transcription and genomic DNA removal were performed using the Hifair^®^ AdvanceFast One-step RT-gDNA Digestion SuperMix for qPCR (Yeasen, 11151ES60). QPCR was carried out with 2× Hieff Canace® AdvanceFast PCR Master Mix (With Dye) (Yeasen, 10164ES03) on an Applied Biosystems QuantStudio 5. Primer sequences used in this study are listed in Supplementary table 1. Assays were performed in duplicate and repeated independently at least three times. Gene expression levels were normalized to internal control and analyzed using the comparative 2^^(-ΔΔCt)^ method.

### RNA decay analysis

Cells were treated with actinomycin D (Selleck, S8964) to inhibit transcription. Total RNA was extracted at indicated time points, reverse-transcribed, and quantified by qRT-PCR. RNA abundance was normalized to the initial time point (t = 0). Decay constants were derived by fitting a first-order exponential decay model, and RNA half-lives were calculated as ln(2)/k.

### RNA immunoprecipitation (RIP)

HEK293T cells were lysed in RIP buffer containing protease and RNase inhibitors (TaKaRa, 2313A). Lysates were clarified by centrifugation and quantified. Protein A beads were pre-coupled with target-specific or control IgG antibodies. After reserving a portion of lysate as input, the remainder was incubated with antibody-coupled beads overnight at 4 °C. Immunoprecipitated complexes were divided for parallel analysis: one portion was processed for immunoblotting, while the other was used for RNA extraction, reverse transcription, and qRT-PCR.

### Chromatin immunoprecipitation (ChIP)

Cells at 70–80% confluence were crosslinked with 1% formaldehyde for 10 min and quenched with glycine. After washing and pelleting, cells were lysed, and nuclei were collected, resuspended in shearing buffer, and sonicated to obtain chromatin fragments (200–1500 bp). Chromatin was pre-cleared and incubated with specific antibodies or IgG control overnight at 4 °C with rotation. Immune complexes were captured with Protein A beads, washed, eluted, and crosslinks were reversed. Purified DNA was analyzed by qRT-PCR using primers targeting regions within 2 kb upstream of the transcription start site.

### Immunoblotting (IB)

Cells were washed with ice-cold PBS, lysed on ice in enhanced RIPA buffer (APPLYGEN, C1053+–100) containing PMSF (Sigma-Aldrich, P7626) and Phosphatase Inhibitor Cocktail (APPLYGEN, P1260). Lysates were clarified by centrifugation at 4 °C, and equal amounts of protein were denatured in SDS buffer and resolved by SDS–PAGE. Proteins were transferred to NC membranes (Cytiva, 10600002), blocked in 5% non-fat milk in TBST for 1 h at room temperature, or 5% BSA for phospho-specific blots. Membranes were incubated with primary antibodies overnight at 4 °C, washed with TBST, and incubated with HRP-conjugated secondary antibodies (Biodragon, BF03001 or BF03008) for 1 h. Bands were visualized using chemiluminescence (Biodragon, BF06053).

### Co-immunoprecipitation (co-IP)

Cells were lysed in ice-cold co-IP buffer containing protease inhibitors. Cleared lysates were incubated overnight at 4 °C with specific antibodies or isotype-matched IgG controls, followed by capture with Protein A agarose beads. After washing, bound proteins were eluted and analyzed by immunoblotting.

### Immunofluorescence

Cells grown on glass coverslips were washed with PBS and fixed with 4% PFA for 10–15 min at room temperature. After washing, cells were permeabilized with 0.1–0.3% Triton X-100 (Sigma-Aldrich, T8787) in PBS for 10 min and blocked in 3–5% BSA in PBS for 30–60 min at room temperature. Coverslips were incubated with primary antibodies diluted in blocking buffer for 1–2 h at room temperature or overnight at 4 °C, washed three times with PBS, and then incubated with appropriate fluorophore-conjugated secondary antibodies (Biodragon, BD9010 or BD9276) for 1 h at room temperature in the dark. Nuclei were counterstained with Hoechst 33258. Coverslips were imaged on a confocal fluorescence microscope under identical acquisition settings for compared samples.

### Cellular thermal shift assay (CETSA)

Confluent cells were treated with DMSO or curcumol (50 μM) for 2 h, collected, washed with PBS, and resuspended in PBS containing protease inhibitors. Aliquots (80 μL) were heated at 37–62 °C (5 °C increments) for 3 min, followed by three freeze-thaw cycles. After centrifugation (13000 g, 10 min, 4 °C), supernatants were mixed with loading buffer and stored at −80 °C. Protein stability was assessed by immunoblotting.

### Drug affinity responsive target stability (DARTS)

Cell lysates were prepared by freeze-thaw cycles in PBS with protease inhibitors and clarified by centrifugation. Aliquots (72 μL) were incubated with curcumol (0–1000 μM) or DMSO at room temperature for 1 h, followed by pronase (MCE, HY-114158) digestion (5 μg/mL) for 1 h. Reactions were stopped with loading buffer, boiled, and stored at −80 °C. IGF2BP3 and β-tubulin levels were analyzed by immunoblotting.

### Animals

All mice were maintained under specific-pathogen-free conditions. Animal experiments were approved by the Institutional Review Board of Peking University (approval number: LA2024099).

### Murine peritoneal macrophage induction and isolation

Mice were injected intraperitoneally with 2.5 mL of fluid thioglycollate medium (FTM) (BD, 225650). After three days, mice were euthanized, and peritoneal lavage was performed using 5 mL of ice-cold PBS. Lavage fluid was collected, centrifuged (600 g, 10 min, 4 °C), and cells were resuspended in PBS, filtered through a strainer, and washed again. Cells were finally resuspended in RPMI-1640 medium supplemented with 10% FBS and antibiotics.

### In vivo VSV infection in Igf2bp3 KO mice

*Age- and sex-matched 6–8-week-old C57BL/6J WT and Igf2bp3 knockout mice (Cyagen, S-KO-01949) were injected via tail vein with VSV (5 × 10⁷ PFU/g body weight). At 24 h post-infection, mice were euthanized, and liver, spleen, and lung tissues were collected. Viral RNA levels were quantified by qRT-PCR, and viral titers were determined by plaque assay from tissue homogenates. For histopathological analysis, lungs were fixed in 4% PFA, paraffin-embedded, sectioned, and stained with H&E*.

### Curcumol treatment in Trex1 KO mice

*Trex1* KO mice (3–4 weeks old) were randomized into DMSO or curcumol groups. Curcumol was formulated in a solvent containing 5% DMSO, 40% PEG300 (Selleck, S6704), 5% Tween-80 (Selleck, S6702), and 50% ddH_2_O. *Trex1* KO mice received daily intraperitoneal injections of curcumol (50 μg/g body weight) or DMSO for 14 consecutive days. WT control groups received daily intraperitoneal injections of DMSO for 14 consecutive days. Twenty-four hours after the final dose, organs were harvested for RNA extraction and qRT-PCR analysis of *Ifnb1*, *Il6*, and *Isg15* expression, as well as for histological evaluation using H&E-stained paraffin sections.

### Quantification and statistical analysis

Statistical analyses were performed using GraphPad Prism 8.0. Data are presented as mean ± SD. Differences between two groups were assessed using unpaired *t*-tests; comparisons among multiple groups were analyzed by one-way ANOVA followed by appropriate post-hoc tests (e.g., Tukey). A *P*-value < 0.05 was considered statistically significant.

## Acknowledgments

We thank Dr. Tomas Lindahl for providing *Trex1*^+/−^ mice (C57BL/6 background). We thank Dr. Feng Shao for providing iBMDMs. We thank Dr. Fuping You for the L929-ISRE cell line, Dr. Zhengfan Jiang for the 2fTGH-ISRE cell line, Dr. Hong-Bing Shu for the luciferase reporter plasmids, Dr. Zhijian Chen for Flag-MAVS plasmid and Dr. T.Maniatis for Flag-TBK1 plasmid.This study was supported by the National Natural Science Foundation of China (82271796). Open Project funded by the Key Laboratory of Carcinogenesis and Translational Research, Ministry of Education, grant 2025 Open Project-5, NHC Key Laboratory of Medical Immunology at Peking University.

## Author Contributions

A.Z. and J.Z. designed the study. A.Z. performed experiments with assistance from S.G., R.C.T., X.Y., X.Z., L.Z. and X.Y.S. A.Z. and J.Z. analyzed data and wrote the manuscript. J.Z. supervised the project and acquired funding.

