## Supplementary information for "IGF2BP3 amplifies antiviral innate immunity with implications for autoimmune diseases"

Jun Zhang*

**Author Contributions:** A.Z. and J.Z. designed the study. A.Z. performed experiments with assistance from S.G., R.C.T., X.Y., X.Z., L.Z. and X.Y.S. A.Z. and J.Z. analyzed data and wrote the manuscript. J.Z. supervised the project and acquired funding.

### Supplementary Figure legends

#### Supplementary Figure 1. IGF2BP3 is selectively induced in viral infection datasets and correlates with type I interferon programs, and is upregulated by diverse innate stimuli *in vitro*

**A-M**, Expression of $m^6A$ readers/writers/erasers in the influenza dataset GSE100160 (healthy controls vs influenza patients), including *YTHDF1* (A), *YTHDF2* (B), *YTHDF3* (C), *YTHDC1* (D), *YTHDC2* (E), *HNRNPG* (F), *IGF2BP1* (G), *IGF2BP2* (H), *IGF2BP3* (I), *METTL3* (J), *WTAP* (K), *ALKBH5* (L), and *FTO* (M).

**N-P,** Expression of IGF2BP family members in the COVID-19 dataset GSE217948 (healthy controls vs COVID-19 patients), including IGF2BP1 (N), IGF2BP2 (O), and IGF2BP3 (P).

**Q**, Correlation analyses identifying genes strongly associated with *IGF2BP3* expression in GSE100160.

**R**, GO enrichment analyses of IGF2BP3-correlated genes from GSE100160.

**S**,**T**, qRT-PCR of *IGF2BP3* mRNA levels in THP-1, iBMDM, BMDM, and HeLa cells stimulated with HSV-1 (S) or LPS (T) for 9 h.

**U**, Relative luciferase activities of IFN-β, ISRE, and NF-κB reporters in HeLa cells transfected with empty vector or an IGF2BP3 expression plasmid and infected with HSV-1 for 24 h.

Data are presented as mean ± SD for cellular assays (S–U). Statistical significance was determined by unpaired two-tailed Student's *t* test; exact *P* values are indicated in the panels.

#### Supplementary Figure 2. *IGF2BP3* knockdown impairs SeV-induced antiviral signaling and promotes VSV replication in A549 cells

**A**, IB analysis validating *IGF2BP3* knockdown in A549 cells stably expressing shNC or shIGF2BP3-1/shIGF2BP3-2.

**B,** Relative luciferase activities of IFN-β, ISRE, and NF-κB reporters in shNC and shIGF2BP3-1/shIGF2BP3-2 A549 cells infected with SeV for 24 h.

**C,D**, qRT-PCR of *IFNB1* (C) and *CXCL10* (D) mRNA levels in shNC and shIGF2BP3-1/shIGF2BP3-2 A549 cells at the indicated time points following SeV infection..

**E**, IB analysis of phosphorylated and total TBK1, IRF3, and p65 in shNC and shIGF2BP3-1/shIGF2BP3-2 A549 cells at the indicated time points following SeV infection..

**F-H**, VSV replication in shNC and shIGF2BP3-1/shIGF2BP3-2 A549 cells infected with VSV-GFP was assessed by GFP imaging at 24 h (F) and by qRT-PCR (VSV RNA; G) and IB (VSV-GFP; H) at 0, 12, and 24 h post-infection. Scale bars, 100 μm.

**I**, Schematic of the acute VSV infection model in *Igf2bp3*$^{+/+}$ and *Igf2bp3*$^{-/-}$ mice infected by tail-vein injection ($5 \times 10^7$ pfu/g), with serum and tissues collected at 24 h post-infection.

**J**, IB analysis of phosphorylated and total TBK1, IRF3, and p65 in peritoneal macrophages isolated from *Igf2bp3*$^{+/+}$ and *Igf2bp3*$^{-/-}$ mice at the indicated time points following SeV infection.

Data are presented as mean ± SD. Statistical significance was determined by unpaired two-tailed Student's *t* test; exact *P* values are indicated in the panels. Data in A, E, H, and J are representative of three independent experiments with similar results.

**Supplementary Figure 3. Validation of *METTL3* knockdown, characterization of an m$^6$A-binding–defective IGF2BP3 mutant, and functional rescue by MAVS or TBK1 in *IGF2BP3* KO cells**

**A**, qRT-PCR of *METTL3* mRNA levels in HEK293T cells transfected with shNC, shMETTL3-1, or shMETTL3-2 at 24 h post-transfection.

**B,** Schematic of the IGF2BP3 m$^6$A-binding-defective mutant generated by mutating GXXG motifs within the KH3/KH4 domains to GEEG.

**C**, IB analysis of IGF2BP3 protein levels in the corresponding samples shown in Figure 3A.

**D**, IB analysis of IGF2BP3 protein levels in the corresponding samples shown in Figure 3B.

**E**, IB analysis of IGF2BP3 protein levels in the corresponding samples shown in Figure 3,C and D.

**F**, IB analysis of IGF2BP3 protein levels in the corresponding samples shown in Figure 3,F and G.

**G**, M$^6$A-RIP–qRT-PCR of *TBK1* mRNA enrichment in HEK293T cells expressing shNC, shMETTL3-1, or shMETTL3-2 using IgG or anti-m$^6$A antibody.

**H**, Flag-RIP–qRT-PCR of *TBK1* mRNA association with Flag–IGF2BP3 WT or Flag–IGF2BP3 mutant in *IGF2BP3* KO HeLa cells reconstituted with Flag–IGF2BP3 WT or Flag–IGF2BP3 mutant using IgG or anti-Flag antibody.

**I**, IB analysis of IGF2BP3 protein levels in the corresponding samples shown in Figure 3I.

Data are presented as mean ± SD. Statistical significance was determined by unpaired two-tailed Student's *t* test; exact *P* values are indicated in the panels. Data in C, D, E, F, and I are representative of three independent experiments with similar results.

**Supplementary Figure 4. Construction of MAVS m$^6$A-site WT/MUT reporters and specificity controls for eIF4G1/RIG-I RIP assays**

**A**, Schematic of pGL3-promoter–MAVS 3′UTR WT and m$^6$A-site-mutant reporters containing a 117-nt *MAVS* fragment with three predicted m$^6$A sites.

**B**, Schematic representation and IGV tracks showing m$^6$A and IGF2BP3-binding peaks across the *TBK1* transcript, based on MeRIP-seq (GSE29714) and IGF2BP3 RIP-seq (GSE261200) datasets.

**C,** Endogenous co-IP of IGF2BP3 association with eIF4G1 in HEK293T cells.

**D-F**, RIP–qRT-PCR of co-precipitated *GAPDH* mRNA (negative control) using IgG or anti-eIF4G1 antibody in A549 cells stably expressing shNC or shIGF2BP3-1/shIGF2BP3-2 (D), in WT and *IGF2BP3* KO HeLa cells (E), and in HEK293T cells transfected with vector or an IGF2BP3 expression plasmid (F), with IB analysis of eIF4G1 protein levels in the corresponding samples shown in Figure 4, M-R and Supplementary Figure 4, D-F.

**G,** RIP–qRT-PCR of co-precipitated *GAPDH* mRNA (negative control) using IgG or anti-RIG-I antibody in WT/*IGF2BP3* KO/reconstituted HeLa cells, with IB analysis of RIG-I protein levels in the corresponding samples shown in Figure 5G and Supplementary Figure 4G.

**H**, IB analysis of IGF2BP3 protein levels in the corresponding samples shown in Figure 5G and Supplementary Figure 4G.

**I**, RIP–qRT-PCR of co-precipitated *GAPDH* mRNA (negative control) using IgG or anti-RIG-I antibody in HEK293T cells transfected with empty vector or Flag–IGF2BP3, with IB analysis of RIG-I protein levels in the corresponding samples shown in Figure 5H and Supplementary Figure 4I.

**J**, IB analysis of Flag-IGF2BP3 protein levels in the corresponding samples shown in Figure 5H and Supplementary Figure 4H.

**K**, IB analysis of IGF2BP3 protein levels in the corresponding samples shown in Figure 5K.

**L**, IB analysis of RIG-I protein levels in the corresponding samples shown in Figure 5K.

Data are presented as mean ± SD. Statistical significance was determined by unpaired two-tailed Student's *t* test; exact *P* values are indicated in the panels. Data in D-K are representative of three independent experiments with similar results.

**Supplementary Figure 5. Positive-control validation for curcumol effects on IGF2BP3 binding, translation engagement, and mRNA stability, and expression/correlation in SLE cohorts**

**A-D**, Expression of IGF2BP family members in SLE cohorts using GSE61635 (*IGF2BP1*, A; *IGF2BP2*, B) and GSE49454 (*IGF2BP1*, C; *IGF2BP2*, D).

**E,** Correlation analyses identifying genes strongly associated with *IGF2BP3* expression in GSE61635.

**F**, GO enrichment analyses of IGF2BP3-correlated genes from GSE61635.

**G**, RIP–qRT-PCR of IGF2BP3-associated *MYC* mRNA (positive control) in HEK293T cells treated with DMSO or curcumol (50 μM) for 24 h using IgG or anti-IGF2BP3 antibody.

**H**, IB analysis of IGF2BP3 protein levels in the corresponding samples shown in Figure 6, G and H and Supplementary Figure 5G.

**I**,**J**, ActD-chase qRT-PCR of *MYC* mRNA decay in HeLa cells (I) or *IGF2BP3* KO HeLa cells (J) pretreated with DMSO or curcumol (50 μM) for 24 h and harvested at 0, 6, 12, and 24 h after ActD treatment (4 μg/mL).

**K**, RIP–qRT-PCR of eIF4G1-associated *MYC* mRNA (positive control) in WT and *IGF2BP3* KO HeLa cells treated with DMSO or curcumol (50 μM) for 24 h using IgG or anti-eIF4G1 antibody.

**L**, IB analysis of eIF4G1 protein levels in the corresponding samples shown in Figure 6, M and N and Supplementary Figure 5K.

Data are presented as mean ± SD. Statistical significance was determined by unpaired two-tailed Student's *t* test; exact *P* values are indicated in the panels. Data in H, and L are representative of three independent experiments with similar results.

**Supplemental Table 1. The Sequences of Primers**

| Primer name | Sequences | |
|---|---|---|
| Human *IFNB1* | Forward | ACTGCCTCAAGGACAGGATG |
| | Reverse | GGCCTTCAGGTAATGCAGAA |
| Human *GAPDH* | Forward | ACCCACTCCTCCACCTTTGA |
| | Reverse | CTGTTGCTGTAGCCAAATTCGT |
| Human *ISG15* | Forward | CGCAGATCACCCAGAAGATCG |
| | Reverse | TTCGTCGCATTTGTCCACCA |
| Human *IGF2BP3* | Forward | TATATCGGAAACCTCAGCGAGA |
| | Reverse | GGACCGAGTGCTCAACTTCT |
| Human *MAVS* | Forward | CAGGCCGAGCCTATCATCTG |
| | Reverse | GGGCTTTGAGCTAGTTGGCA |
| Human *TBK1* | Forward | TGGGTGGAATGAATCATCTACGA |
| | Reverse | GCTGCACCAAAATCTGTGAGT |
| Human *METTL3* | Forward | TTGTCTCCAACCTTCCGTAGT |
| | Reverse | CCAGATCAGAGAGGTGGTGTAG |
| Human *CXCL10* | Forward | CTCTAAGTGGCATTCAAGGA |
| | Reverse | GGATTCAGACATCTCTTCTCA |
| Human *MYC* | Forward | TTCGGGTAGTGGAAAACCAG |
| | Reverse | AGTAGAAATACGGCTGCACC |
| IGF2BP3-Promoter-Site1 | Forward | CCACTAGAACCCCAGCACTG |
| | Reverse | TACAATGGTGGTTGTCGGGG |
| IGF2BP3-Promoter-Site2 | Forward | GCCTTTTCTGAATCGCGCAA |
| | Reverse | TTCTAAGGCTGGCATGGACG |
| IGF2BP3-Promoter-Site3 | Forward | TTCCTCCTCCCCCTTCTCAG |
| | Reverse | CGTCTCACGTGAGGAATCCC |
| Mouse *Ifnb1* | Forward | CAGCTCCAAGAAAGGACGAAC |
| | Reverse | GGCAGTGTAACTCTTCTGCAT |
| Mouse *Ifna4* | Forward | GTTCCAGAAGGCTCAAGCCATC |
| | Reverse | TAGGAGGCTCTGTTCCCAAGCA |
| Mouse *Cxcl10* | Forward | GCCGTCATTTTCTGCCTCA |
| | Reverse | CGTCCTTGCGAGAGGGATC |
| Mouse *Il6* | Forward | TACCACTTCACAAGTCGGAGGC |
| | Reverse | CTGCAAGTGCATCATCGTTGTTC |
| Mouse *Isg15* | Forward | TGACTGTGAGAGCAAGCAGC |
| | Reverse | CCCCAGCATCTTCACCTTTA |
| VSV-RNA | Forward | ACGGCGTACTTCCAGATGG |
| | Reverse | CTCGGTTCAAGATCCAGGT |
| Human-METTL3-shRNA1 | | GCCCAAGTGCAAGAATTCTGT |
| Human-METTL3-shRNA2 | | GGGCCCAAGTGCAAGAATTCT |
| IGF2BP3-423/424 | Forward | GGTCCCCAAAGGGATGAGAA |
| | Reverse | TGAGGGTCTGGGCCATAGAA |
| IGF2BP3-505/506 | Forward | TTTGCTGCTGGCAGAGTTATTGGAGAGGAAGGCAAAACGGTGAATG |
| | Reverse | CATTCACCGTTTTGCCTTCCTCTCCAATAACTCTGCCAGCAGCAAA |

| | | |
|---|---|---|
| pGL3-promoter-MAVS-Mut | Forward | GAAAGATCGCCGTGTAATTCTAGAATGCCGTTTGCTGAAGGCAAGGC |
| | Reverse | GGCCGGCCGCCCCGACTCTAGACTGGCCTCTTGCTGTG |
| pGL3-promoter-MAVS-WT | Forward | GAAAGATCGCCGTGTAATTCTAGAATGCCGTTTGCTGAAG |
| | Reverse | GGCCGGCCGCCCCGACTCTAGACTGGTCTCTTGCTGTG |
| Human-IGF2BP3-sgRNA | | CTTACCTCCCACTGTAAATG |
| Mouse-IGF2BP3-sgRNA | | ACTCGGTCCCTAAACGGCAG |
| Human-IGF2BP3-shRNA1 | | GAAACTTCAGATACGAAATAT |
| Human-IGF2BP3-shRNA2 | | AATCGATGTCCACCGTAAAGA |
