## Supplementary figures and images for "IGF2BP3 amplifies antiviral innate immunity with implications for autoimmune diseases"

### Supplementary Figure 1

**Supplementary Figure 1**

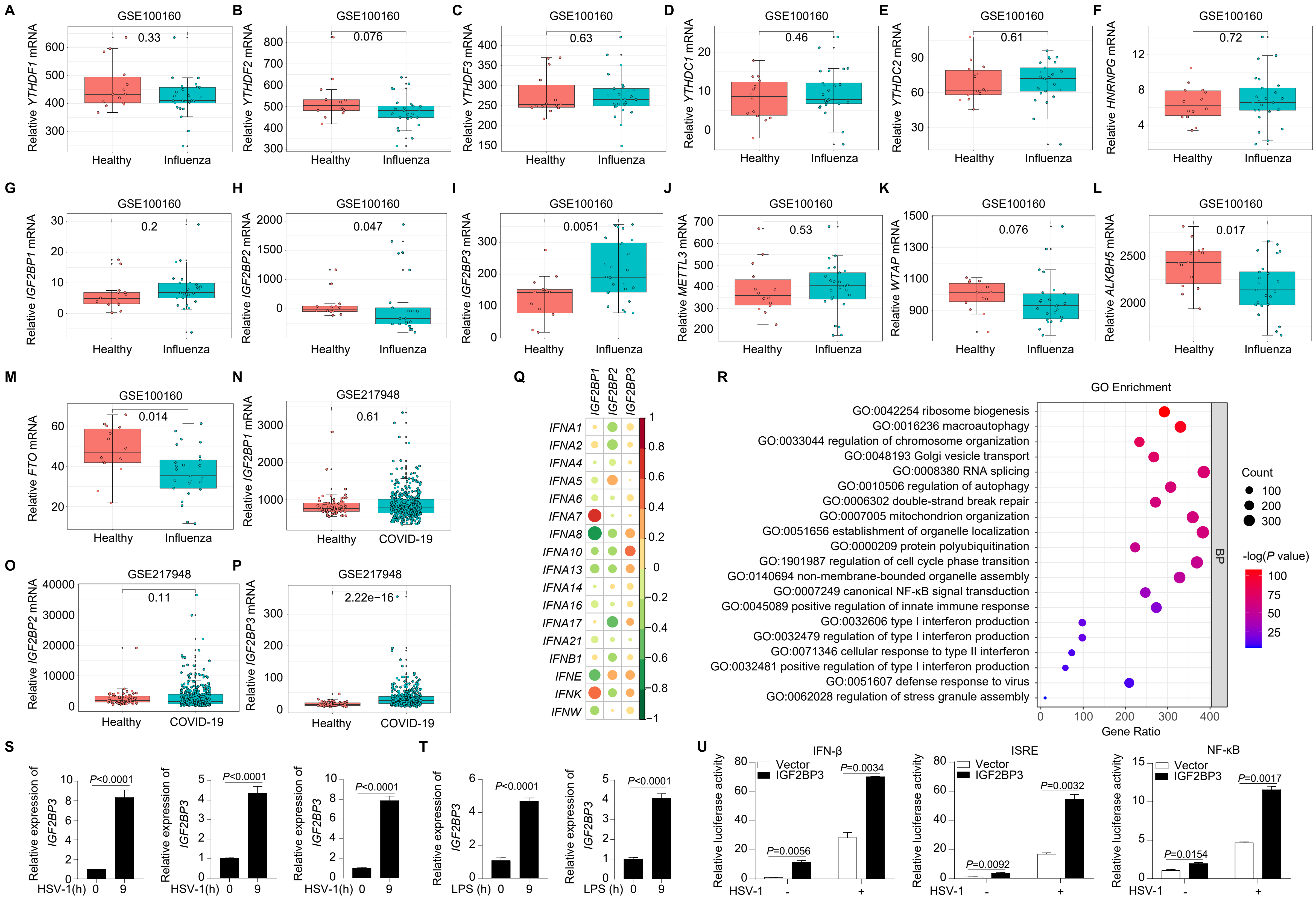

### Supplementary Figure 2

Supplementary Figure 2

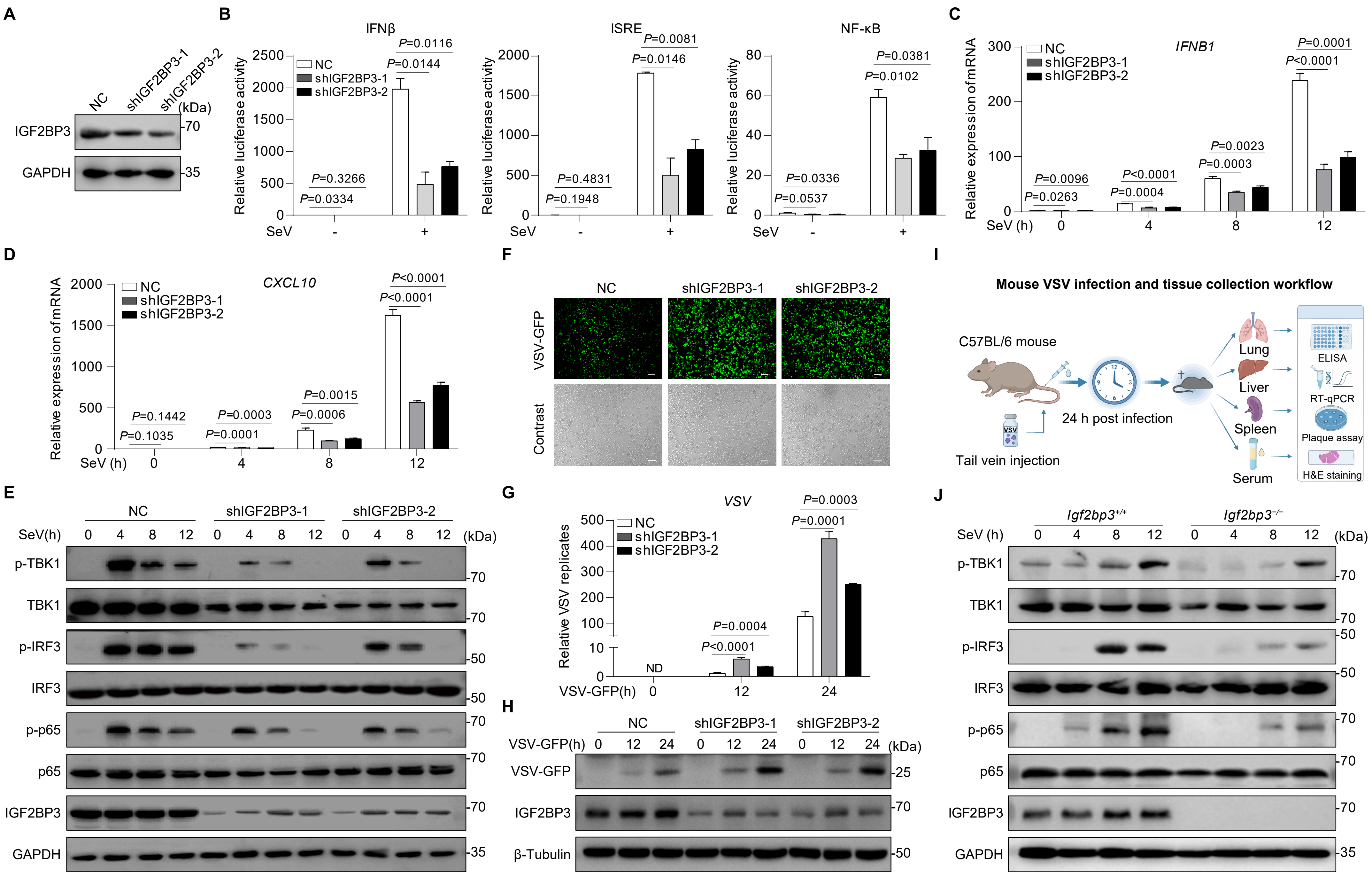

### Supplementary Figure 3

Supplementary Figure 3

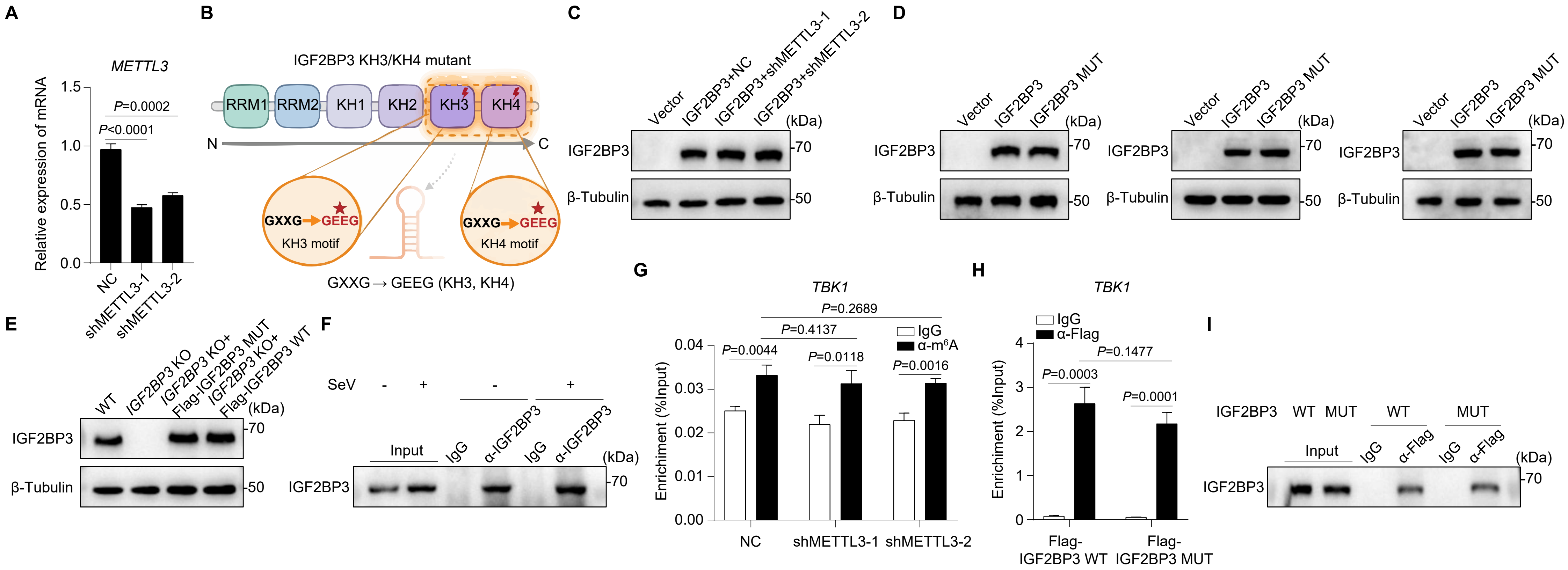

### Supplementary Figure 4

Supplementary Figure 4

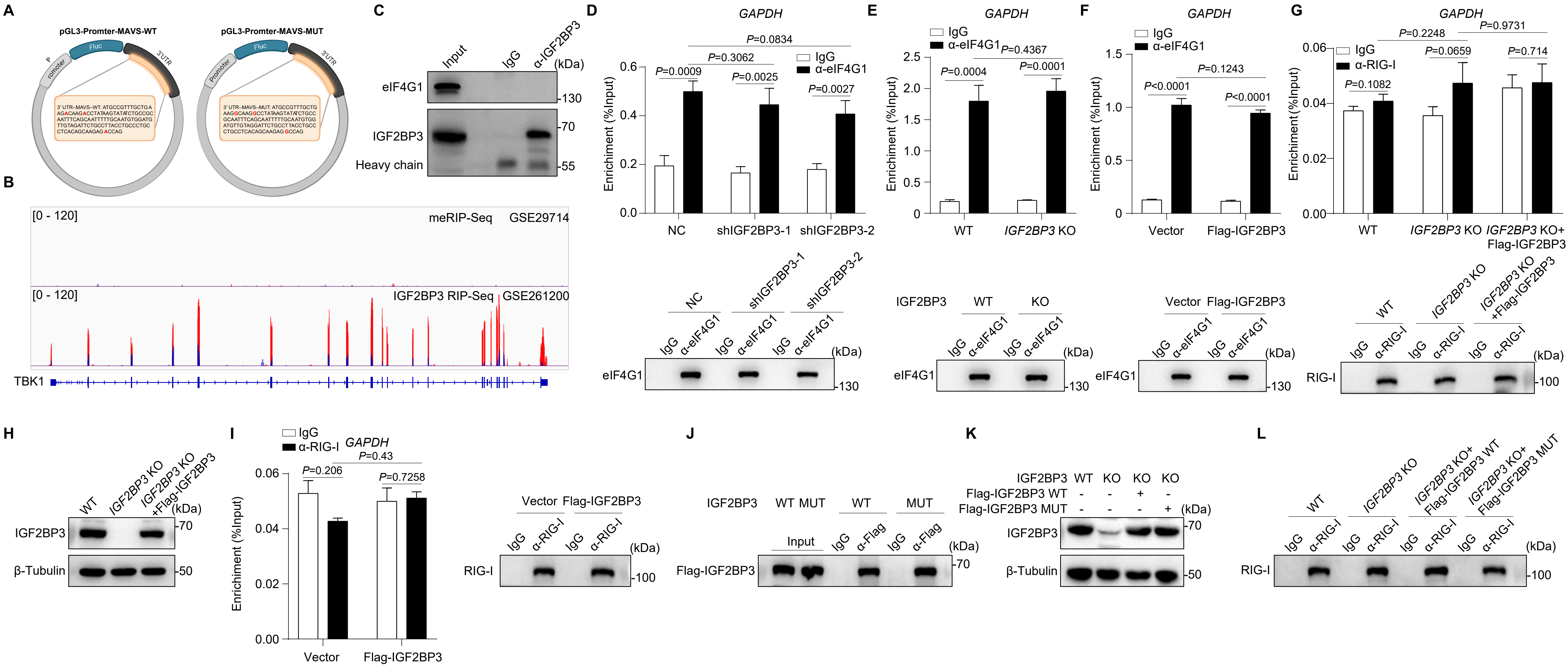

### Supplementary Figure 5

**Supplementary Figure 5**

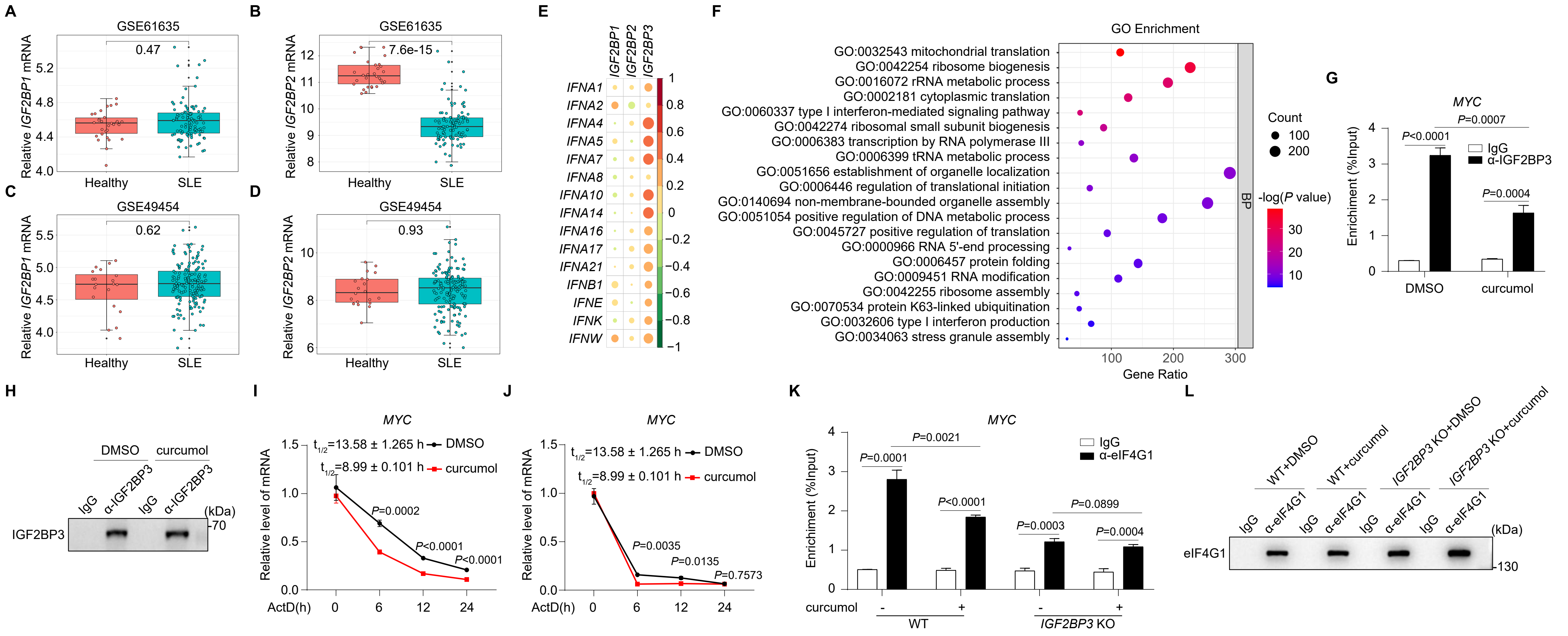
